# 5’Untranslated regions provide a versatile toolkit for tunable exogenous protein expression

**DOI:** 10.64898/2026.01.19.700389

**Authors:** Camila Garcia, Dylan Poch, Aden M. Alemayhu, Candice E. Paulsen

## Abstract

Transient transfection is widely used for protein expression in heterologous systems, yet uncontrolled overexpression frequently introduces artifacts that confound functional analyses. Although stable cell lines can mitigate these issues, generating lines for multiple constructs or variants is often impractical. Common alternatives, such as DNA titration, altered transfection conditions, or promoter swapping, provide only coarse and inconsistent control of protein abundance. Here, we establish a panel of ten human 5’ untranslated regions (5’UTRs) as a modular strategy to tune protein expression during transient transfection. Across three soluble proteins and three membrane proteins, these 5’UTRs produce a reproducible dynamic range of expression, including fine-grained control of eYFP and the large sensory ion channel TRPA1. Notably, one 5’UTR consistently suppresses expression across all proteins tested and alleviates overexpression-associated artifacts, improving functional analysis of a hyperactive channel variant, substantially reducing background in proximity biotinylation assays, and enhancing the specificity of a stress granule marker. In contrast, most 5’UTRs enhance expression of the TRPV1 and TRPM8 sensory receptors, improving protein yield in heterologous systems. Together, this work identifies 5’UTRs as a compact, versatile, and broadly applicable tool to fine-tune protein abundance, enabling more physiologically relevant and assay-optimized expression in transient transfection experiments.

**Significance Statement:**

- Protein overexpression remains a pervasive limitation in transient transfection experiments, as commonly used strategies provide coarse, labor-intensive, and often unpredictable control over protein abundance, frequently leading to artifacts or loss of physiological relevance.
- A compact, modular 5’UTR toolkit enables fine-grained and scalable control of protein abundance in transient transfection, providing a reproducible dynamic range across soluble and membrane proteins outperforming promoter swapping for continuous tuning of expression levels.
- Tunable expression improves experimental fidelity across diverse applications, allowing investigators to match protein abundance to specific assay needs, reducing overexpression artifacts, preserving physiological readouts, and enhancing yield when high expression is required.

## INTRODUCTION

Protein overexpression occurs when a protein is produced at levels exceeding those that are physiologically relevant, often leading to experimental artifacts and misinterpretation of function (Bolognesi and Lehner 2018; Moriya 2015). Heterologous protein expression has traditionally relied on both stable and transient transfection approaches (Kim and Eberwine 2010). In stable transfection, foreign genetic material is integrated into the host genome, enabling constitutive or inducible expression that is maintained across cell divisions (Gray 1997; Recillas-Targa 2006; Chong, Yeap, and Ho 2021). In contrast, transient transfection introduces plasmid DNA or mRNA that is expressed episomally without genomic integration, allowing rapid and flexible protein production for small-scale assays and method development (Chong, Yeap, and Ho 2021; Dalton and Barton 2014; Alkharobi 2023). Although both approaches can result in overexpression, transient transfection is particularly prone to excessive and poorly controlled protein levels due to variable expression kinetics and plasmid copy number (Makrides 2003; Fus-Kujawa et al. 2021).

In certain applications – such as protein purification or structural studies – high expression levels are advantageous or even required (Ušaj et al. 2018). However, many experimental questions demand intermediate or low protein abundance, including studies of protein regulation, trafficking, turnover, proteomics, electrophysiology, and signaling (Di Blasi et al. 2021). In these contexts, overexpression can perturb cellular pathways, alter subcellular localization, or induce cytotoxicity, underscoring the need for modular strategies that enable precise control of protein abundance. Regulatory elements acting at both the transcriptional and translational levels offer potential solutions. Eukaryotic promoters control transcription initiation by recruiting RNA polymerase II and associated factors, and promoter architecture – including motif composition – strongly influences expression output (Conaway et al. 1991; Einarsson et al. 2022; Moritz, Becker, and Göpfert 2015). In contrast, 5’ untranslated regions (5’UTRs) regulate translation by modulating mRNA stability, ribosome recruitment, and translational efficiency, thereby shaping protein output independently of transcription (Hinnebusch, Ivanov, and Sonenberg 2016; Wieder et al. 2025; Ryczek, Łyś, and Makałowska 2023).

Recently, a systematic screen of approximately 30,000 human 5’UTRs identified ten sequences with distinct effects on ribosome recruitment and translational efficiency (Lewis et al. 2025). Here, we validate and extend the use of this 5’UTR panel as a strategy to tune protein expression in transient transfection systems, directly addressing overexpression-associated artifacts. We focus primarily on three members of the Transient Receptor Potential (TRP) ion channel family – TRPA1, TRPV1, and TRPM8 – which are homotetrameric, calcium (Ca^2+^)-permeable channels implicated in pain and inflammation that are widely studied in heterologous expression systems (Wang et al. 2008; Julius 2013; Bamps et al. 2021; Bali et al. 2023; Kwon et al. 2021; Sanders et al. 2025; Yin et al. 2024). For these channels, lower expression levels are often required for physiologically meaningful functional measurements, whereas higher expression is advantageous for structural and biochemical analyses, making context-dependent and tunable control of protein abundance particularly valuable.

Our results demonstrate that these 5’UTRs enable a 10-fold dynamic range of protein expression for both the soluble protein eYFP and the large TRPA1 ion channel, while predominantly enhancing expression of TRPV1 and TRPM8. In direct comparison, promoter swapping yielded more binary, high-or-low expression outcomes with limited tunability. We further show that the 5’UTRs can be leveraged to adjust protein abundance across a range of experimental contexts. These include enhancing protein production for purification pipelines, suppressing expression for improved assay sensitivity, and improving the physiological relevance of a heterologously expressed stress granule biomarker associated with neurodegenerative disorders. Together, this work establishes 5’UTRs as a versatile and modular tool for fine-tuning protein expression in transient transfection, expanding the experimental toolkit for achieving physiologically relevant and assay-optimized protein levels across a broad range of biological applications.

## RESULTS

### Eukaryotic promoters function as high-low expression switches

A common strategy to modulate ectopic protein expression is to select promoters with defined transcriptional strength (Forrest et al. 2014; Qin et al. 2010). Eukaryotic promoters serve as platforms for RNA polymerase II and transcriptional co-factors that initiate mRNA synthesis (Conaway et al. 1991). The cytomegalovirus (CMV) promoter is widely used to drive strong expression (Bäck et al. 2019), whereas promoters such as SV40 yield comparatively lower expression in heterologous systems (Yoon et al. 2024; Qin et al. 2010). To define the expression range achievable across commonly used RNA polymerase II promoters, we compared CMV, SV40, PGK, EF1α and UbC in parallel.

We first constructed plasmids encoding 3xFLAG-eYFP under each promoter (**Figures 1A** and **2A**) and transiently transfected them into HEK293T cells. Quantification by fluorescence measurements and anti-FLAG Western blot showed that these promoters segregate into two classes: CMV and EF1α produced high eYFP expression, whereas PGK, SV40, and UbC generated similarly low expression (**Figures 2B-D**). Across promoters, protein abundance varied ∼3-fold by Western blot and ∼10-fold by fluorescence (**Figures 2B** and **D**). The wider range with the fluorescent plate reader measurements than the anti-FLAG expression measurements may speak to the relative sensitives of these readouts.

**Figure 1.**
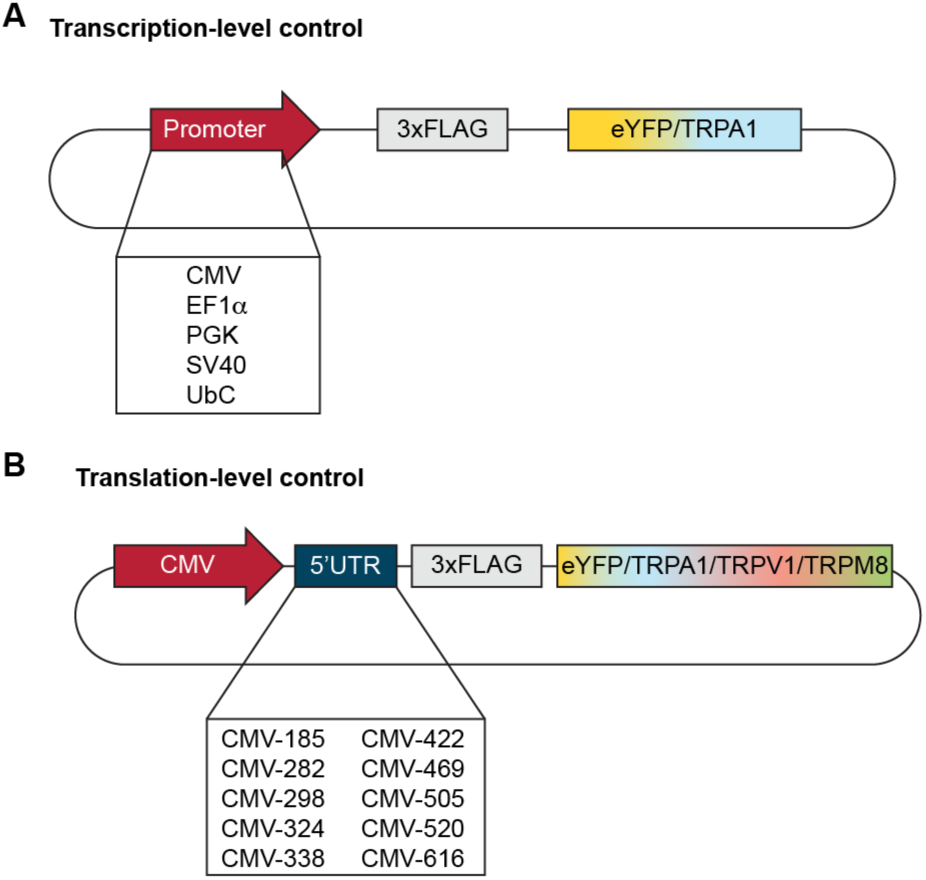
Eukaryotic promoters and human 5’untranslated regions to control exogenous protein expression. (**A**) Cartoon depiction of constructs made with five eukaryotic promoters (red), a 3xFLAG tag (gray), followed by the protein of interest (eYFP or hTRPA1). (**B**) Cartoon depiction of constructs made with a CMV promoter (red) followed by ten 5’untranslated regions (blue), a 3xFLAG tag (gray), and the protein of interest (eYFP, human TRPA1, human TRPV1, or mouse TRPM8).

**Figure 2.**
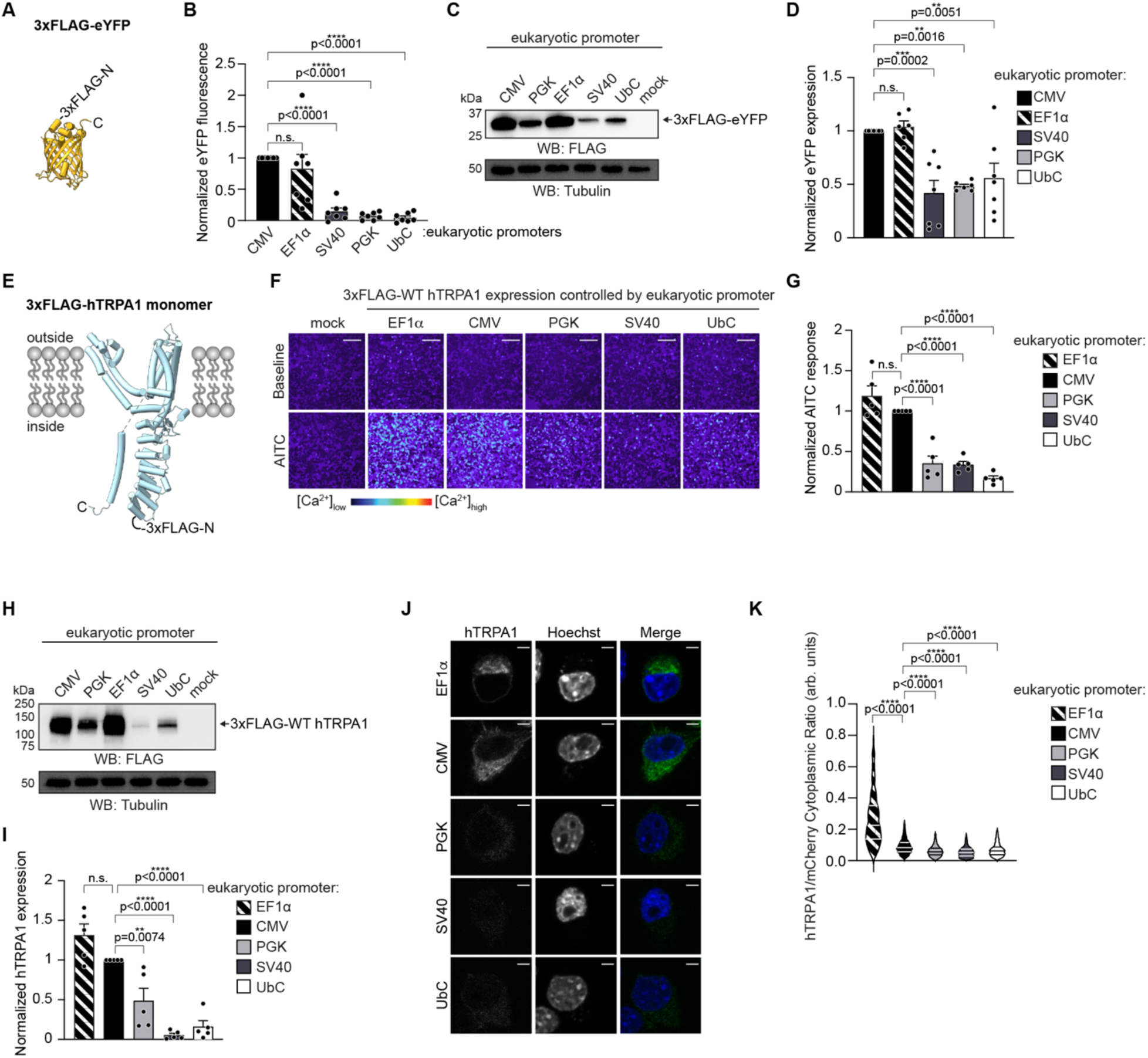
Eukaryotic promoters function as high-low expression switches. (A) Ribbon diagram of the eYFP atomic model (PDB: 3V3D); each monomer is 30.4 kDa. (**B**) Quantification of CMV-normalized eYFP fluorescence for CMV (black), EF1α (black stripes), SV40 (dark gray), PGK (gray), or UbC (white) 3xFLAG-tagged eYFP promoter variants. (**C**) Western blot of lysates from cells expressing 3xFLAG-tagged eYFP controlled by the indicated promoters, probed using HRP-conjugated anti-FLAG antibody. Tubulin was the loading control. Results were verified in seven independent experiments. (**D**) Quantification of CMV-normalized expression of eYFP promoter constructs from data as in (C). (**E**) Monomeric ribbon diagram of the human TRPA1 atomic model for residues K446-E1079 (PDB: 6PQQ); each monomer is 130.5 kDa. (**F**) Ratiometric calcium imaging of HEK293T cells transfected with empty vector (mock), or 3xFLAG-hTRPA1 controlled by the indicated eukaryotic promoters. Cells were stimulated with 100 µM AITC. Images are representative of five independent experiments and are arranged from highest-to-lowest activity. Scale bars indicate 100 µm. (**G**) Quantification of CMV-normalized AITC-evoked change in Fura-2 ratio for 3xFLAG-hTRPA1 controlled by the indicated eukaryotic promoters. Data represent an average of *n* ≥ 2,305 cells per transfection condition per biological replicate. (**H**) Western blot of lysates from cells expressing 3xFLAG-hTRPA1 controlled by the indicated eukaryotic promoters, probed using HRP-conjugated anti-FLAG antibody. Tubulin was the loading control. Results were verified in five independent experiments. (**I**) Quantification of the CMV-normalized expression of 3xFLAG-hTRPA1 controlled by the indicated promoters from data as in (H). (**J**) Representative deconvoluted images of Neuro2A cells expressing 3xFLAG-hTRPA1 controlled by the indicated eukaryotic promoters. Cells were stained with Hoechst (blue) and anti-TRPA1 (green) antibody. Scale bar indicates 4 µm. (**K**) Violin plot quantification of N2A cells co-expressing the indicated 3xFLAG-hTRPA1 constructs and free mCherry plotted as hTRPA1/mCherry cytoplasmic expression ratio (arbitrary units, arb. units). Data represent mean ± SEM. Each condition was tested with an *n* ≥ 500 cells per transfection condition. P-values listed in figure panel, one-way ANOVA with Bonferroni’s *post hoc* analysis. (B, D) Data represent mean ± SEM. *n* = 7 independent experiments, P-values listed in figure panel, one-way ANOVA with Bonferroni’s *post hoc* analysis. (G, I) *n* = 5 independent experiments, P-values listed in figure panel, one-way ANOVA with Bonferroni’s *post hoc* analysis.

To determine whether this behavior extended to larger membrane proteins, we replaced eYFP with the 130.5 kDa human TRPA1 (hTRPA1) ion channel (**Figures 1A** and **2E**). hTRPA1 is a calcium (Ca^2+^)-permeable, non-selective cation channel expressed in peripheral sensory neurons of the pain pathway (Julius 2013). HEK293T cells expressing each promoter construct exhibited functional hTRPA1 activity as assessed by ratiometric Ca^2+^ imaging with Fura-2 and upon allyl isothiocyanate (AITC) stimulation (**Figures 2F**). As observed with eYFP, CMV and EF1α drove higher channel activity and expression, while PGK, SV40, and UbC drove lower levels – a ∼2-fold range in activity and ∼10-fold in expression (**Figures 2F**-**I**).

We next evaluated the hTRPA1 constructs in Neuro2A (N2A) cells, a more physiologically relevant neuronal background (Julius 2013). Immunofluorescence (IF) quantification again revealed a high-or-low pattern: EF1α and CMV yielded the highest hTRPA1 expression, whereas PGK, SV40, and UbC produced minimal signal (**Figures 2J-K** and **S1A**). As in HEK293T cells, these expression levels spanned an ∼10-fold range. Notably, EF1α drove significantly higher expression than CMV in N2A cells, demonstrating cell type-dependent promoter performance (**Figures 2K**).

Together, these findings indicate that PGK, SV40, and UbC consistently confer low expression, while CMV and EF1α confer high expression across both small and large proteins and in multiple cell backgrounds. These promoters, therefore, function effectively as high-low expression switches, offering limited resolution for intermediate expression levels.

### 5’UTRs provide graded control of eYFP expression

5’ untranslated regions (5’UTRs) offer an alternative strategy to modulate ectopic protein expression (Wieder et al. 2025). Whereas promoters regulate transcription at the DNA level, 5’UTRs act post-transcriptionally by influencing translation initiation efficiency and mRNA stability, thereby adjusting the amount of protein produced (Hinnebusch, Ivanov, and Sonenberg 2016). Prior studies identified a panel of ten 5’UTRs capable of tuning expression of a small GFP-nanoLuciferase protein across a ∼10-fold range under a CMV promoter in HEK293T cells (Lewis et al. 2025). Notably, this control was graded rather than the discrete high-or-low pattern observed with promoters.

To test whether this panel behaved similarly in our system, we inserted each 5’UTR individually between the CMV promoter and the 3xFLAG-eYFP start codon (**Figures 1B** and **S2**). The starting plasmid lacking a 5’UTR (CMV only) is referred to as “reference plasmid” throughout the rest of this study. Because 5’UTRs are markedly shorter than promoters (∼600 versus <170 nucleotides), their cloning was straightforward. Transient transfection into HEK293T cells followed by eYFP fluorescence measurements and anti-FLAG Western blot revealed graded expression spanning ∼10-20-fold (**Figures 3A-C** and **S3A**). For example, relative to the reference construct, CMV-469 produced similar expression, CMV-422 yielded ∼2-fold lower, CMV-338 ∼5-fold lower, and CMV-520 ∼18-fold lower fluorescence (**Figure 3A**). Notably, this ranked order of effects agrees with previous work on GFP-nanoLuciferase (Lewis et al. 2025). Thus, 5’UTRs modulate expression along a continuum rather than functioning as binary expression switches.

**Figure 3.**
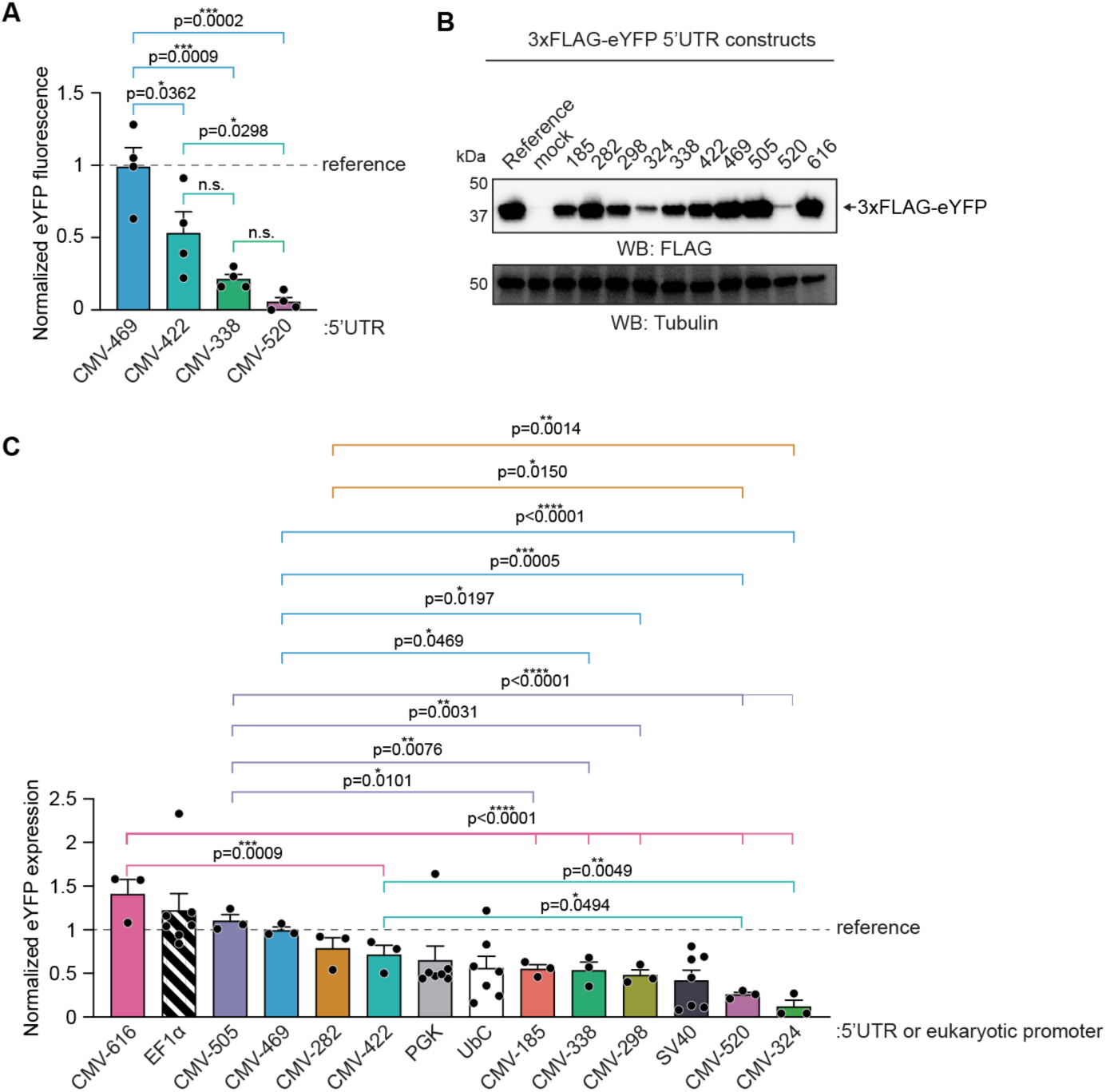
5’UTRs provide graded control of eYFP expression. (**A**) Reference plasmid (CMV promoter, no 5’UTR)-normalized eYFP fluorescence quantification of 3xFLAG-eYFP controlled by CMV-469, CMV-422, CMV-338, or CMV-520. Data represent mean ± SEM. *n* = 4 independent experiments, P-values listed in figure panel, one-way ANOVA with Tukey’s *post hoc* analysis. (**B**) Western blot of lysates from cells expressing 3xFLAG-tagged eYFP variants from experiments as in (A), probed using HRP-conjugated anti-FLAG antibody. Tubulin was the loading control. Results were verified in three independent experiments. (**C**) Quantification of the reference plasmid-normalized expression of 3xFLAG-eYFP controlled by CMV with the indicated 5’UTRs from data as in (B). Promoter data from Figure 2D are replotted for comparison. *n* = 7 (EF1α, SV40, PGK, or UbC) or *n* = 3 (5’UTR constructs) independent experiments. P-values listed in figure panels, one-way ANOVA with Tukey’s *post hoc* analysis among the 5’UTR samples.

### 5’UTRs tune hTRPA1 expression across low, intermediate, and high levels

We next evaluated whether this graded control extended to a larger, multi-pass membrane protein. We replaced eYFP with the hTRPA1 open reading frame in the same 3xFLAG-5’UTR constructs (**Figure 1B**), preserving a common N-terminal context for direct comparison. Following transient transfection into HEK293T cells, Ca^2+^ imaging confirmed functional channels with all 5’UTRs (**Figure 4A**). Chanel activity varied ∼2-fold – from lowest with CMV-520 to intermediate with CMV-185 and similar-to-reference activity with CMV-505 (**Figures 4B** and **S3B**). Western blot revealed a broader ∼10-fold range of protein abundance, with CMV-520 driving the lowest expression, CMV-338 intermediate, and CMV-505 among the highest (**Figures 4C** and **4D**). Although Ca^2+^ imaging reflected the overall 5’UTR ranking, the functional dynamic range was narrower than that observed by Western blot or for eYFP. Transfection efficiency did not account for these differences in expression observed with the 5’UTRs (**Figure S3C**).

**Figure 4.**
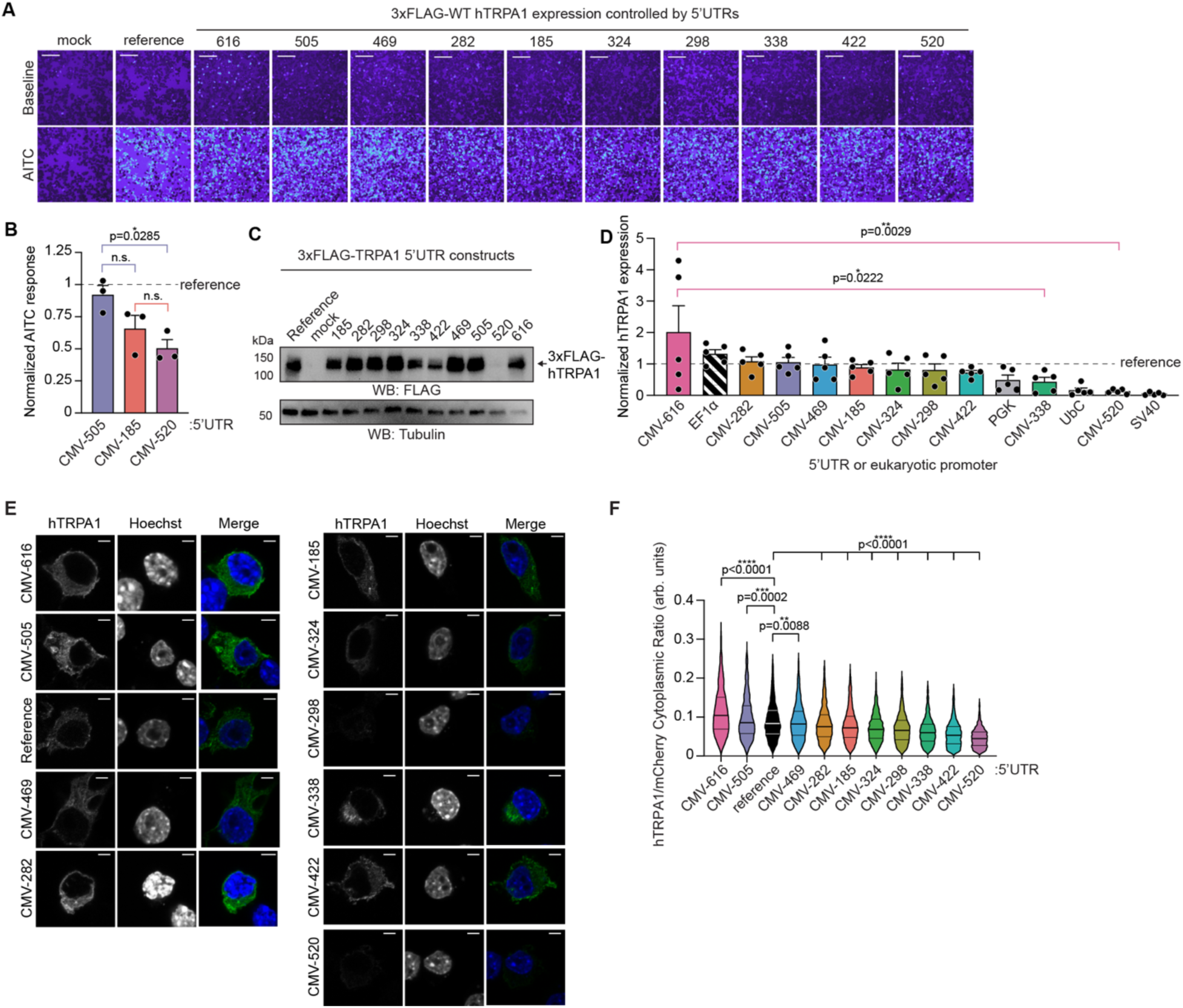
5’UTRs tune hTRPA1 expression across low, intermediate, and high levels. (**A**) Ratiometric calcium imaging of HEK293T cells transfected with empty vector (mock) or 3xFLAG-hTRPA1 controlled by a CMV promoter alone (reference) or with the indicated 5’UTR. Cells were stimulated with 100 µM AITC. Images are representative of three independent experiments and are organized from highest to lowest expression (see panel D). Scale bars indicate 100 µm. (**B**) Quantification of reference plasmid-normalized AITC-evoked changes in Fura-2 ratio of 3xFLAG-hTRPA1 controlled by a CMV promoter with CMV-505, CMV-185, and CMV-520. Data represent mean ± SEM. *n* = 3 independent experiments, with an average of *n* ≥ 2,324 cells per transfection condition per biological replicate. P-values listed in figure panel, one-way ANOVA with Tukey’s *post hoc* analysis. (**C**) Western blot of lysates from cells transfected with empty vector (mock), CMV promoter with no 5’UTR (reference), or 5’UTR 3xFLAG-tagged hTRPA1 constructs, probed using HRP-conjugated anti-FLAG antibody. Tubulin was the loading control. Results were verified in three independent experiments. (**D**) Quantification of reference plasmid-normalized expression of 5’UTR 3xFLAG-tagged hTRPA1 constructs compared to hTRPA1 promoter constructs from data as in (C). Promoter data from Figure 2I are replotted for comparison. *n* = 5 independent replicates. P-values listed in figure panel, one-way ANOVA with Tukey’s *post hoc* analysis among the 5’UTR samples. (**E**) Representative deconvoluted images of Neuro2A cells expressing 3xFLAG-hTRPA1 controlled by a CMV promoter without (reference) or with the indicated 5’UTRs. Cells were stained with Hoechst (blue) and anti-TRPA1 (green) antibody. Scale bar indicates 4 µm. (**F**) Violin plot quantification of N2A cells co-expressing the indicated 3xFLAG-hTRPA1 constructs and free mCherry plotted as hTRPA1/mCherry cytoplasmic expression ratio (arbitrary units, arb. units). Data represent mean ± SEM. Each condition was tested with an *n* ≥ 500 cells per transfection condition. P-values listed in figure panel, one-way ANOVA with Bonferroni’s *post hoc* analysis.

We then assessed hTRPA1 expression in N2A cells by IF imaging and Western blot. As in HEK293T cells, CMV-616 and CMV-505 produced the highest expression, CMV-185, CMV-324, and CMV-298 yielded intermediate levels, and CMV-520 generated minimal expression (**Figures 4E, 4F**, **S1B**). Across these extremes, hTRPA1 expression in N2A cells spanned ∼3-fold by IF imaging and ∼13-fold by Western blot.

Together, these results demonstrate that 5’UTRs tune expression of both small and large ectopic proteins in a graded, multi-level manner rather than the high-low switch-like behavior observed with promoters. Although the precise rank order varies between proteins, consistent grouping into high, intermediate, and low expression drivers underscores the utility of 5’UTRs as versatile tools for fine control of protein expression in transiently transfected cells.

### CMV-520 enables improved detection of a hyperactive hTRPA1 variant

Ratiometric Ca^2+^ imaging is widely used to screen for hyperactive ion channel mutants, but strong overexpression from the CMV promoter can saturate the Fura-2 signals, masking functional differences (Bali et al. 2023). In addition, hyperactive TRPA1 mutants can cause Ca^2+^ overload and cytotoxicity, complicating detailed electrophysiological characterization. We therefore asked whether CMV-520 – a 5’UTR that drives the lowest hTRPA1 expression – could enhance functional detection of the N855S hyperactive variant associated with familial episodic pain syndrome (**Figure 5A**) (Kremeyer et al. 2010). Although N855S exhibits ∼5-fold increased activity by electrophysiology, its enhanced function is difficult to resolve by Ca^2+^ imaging (Bali et al. 2023; Kremeyer et al. 2010).

**Figure 5.**
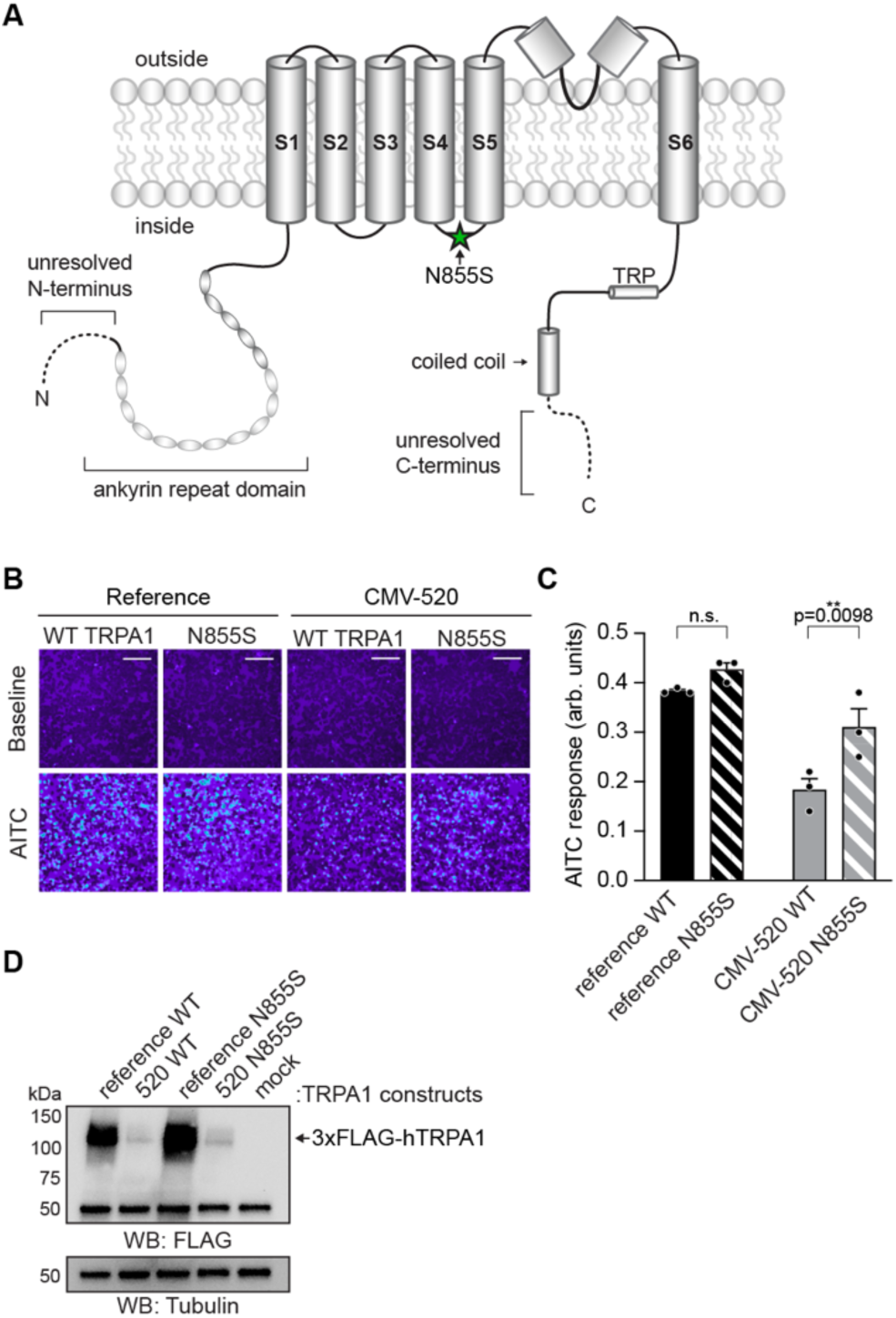
CMV-520 enables improved detection of a hyperactive hTRPA1 variant. (**A**) Cartoon schematic of full-length hTRPA1 monomeric subunit with relevant structural features and a previously identified mutation N855S (green star) denoted. Dashes and transparencies indicate structurally unresolved elements. (**B**) Ratiometric calcium imaging of HEK293T cells transfected with 3xFLAG-wild type (WT) or N855S hTRPA1 controlled by a CMV promoter alone (reference) or with the CMV-520 5’UTR. Cells were stimulated with 100 µM AITC. Images are representative of three independent experiments. Scale bars indicate 100 µm. (**C**) Quantification of AITC-evoked change in Fura-2 ratio (arbitrary units, arb. units) for 3xFLAG-WT or N855S hTRPA1 controlled by a CMV promoter alone (black and black stripe, respectively) or with CMV-520 (grey and grey stripe, respectively). Data represent mean ± SEM. *n* = 3 independent experiments, with an average of *n* ≥ 2,422 cells per transfection condition per biological replicate. P-values listed in figure panel, two-way ANOVA with Bonferroni’s *post hoc* analysis. (**D**) Western blot of lysates from cells expressing 3xFLAG-hTRPA1 variants or transfected with empty vector (mock) from experiments as in (B), probed using HRP-conjugated anti-FLAG antibody. Tubulin was the loading control. Results were verified in three independent experiments.

Consistent with prior work, N855S showed visually increased activity under standard CMV expression, but the difference was not statistically significant (**Figures 5B** and **C**). Incorporating CMV-520 markedly reduced both wild type (WT) and N855S hTRPA1 expression and, in turn, improved the dynamic range of the assay. Under reduced expression, N855S displayed a clear and significant ∼2-fold increase in activity relative to WT hTRPA1 (**Figures 5B-D**).

Thus, lowering channel expression with a weak 5’UTR such as CMV-520 enhances the ability of Ca^2+^ imaging screens to detect hyperactive hTRPA1 variants and may similarly facilitate downstream electrophysiological analyses.

### 5’UTRs as a method to enhance TRPV1 expression and improve protein production

Because the 5’UTRs tuned hTRPA1 across a ∼2-fold functional range and a ∼10-fold expression range, we next tested whether these elements similarly modulate expression and activity of the smaller pain receptor, TRPV1 (**Figure 6A**). Like TRPA1, TRPV1 is a Ca^2+^-permeable non-selective cation channel (Basbaum et al. 2009; Julius 2013). Strikingly, except for CMV-520, all 5’UTRs drove robust hTRPV1 expression in transiently transfected HEK293T cells compared to the reference construct, as measured by anti-FLAG Western blot (**Figures 6B**, **C**, and **S4A**). CMV-324 produced expression comparable to the reference plasmid, whereas the remaining 5’UTRs enhanced expression, enabling a ∼25-fold control of hTRPV1 abundance across high, intermediate, and low levels (**Figures 6B**, **C**, and **S4A**).

**Figure 6.**
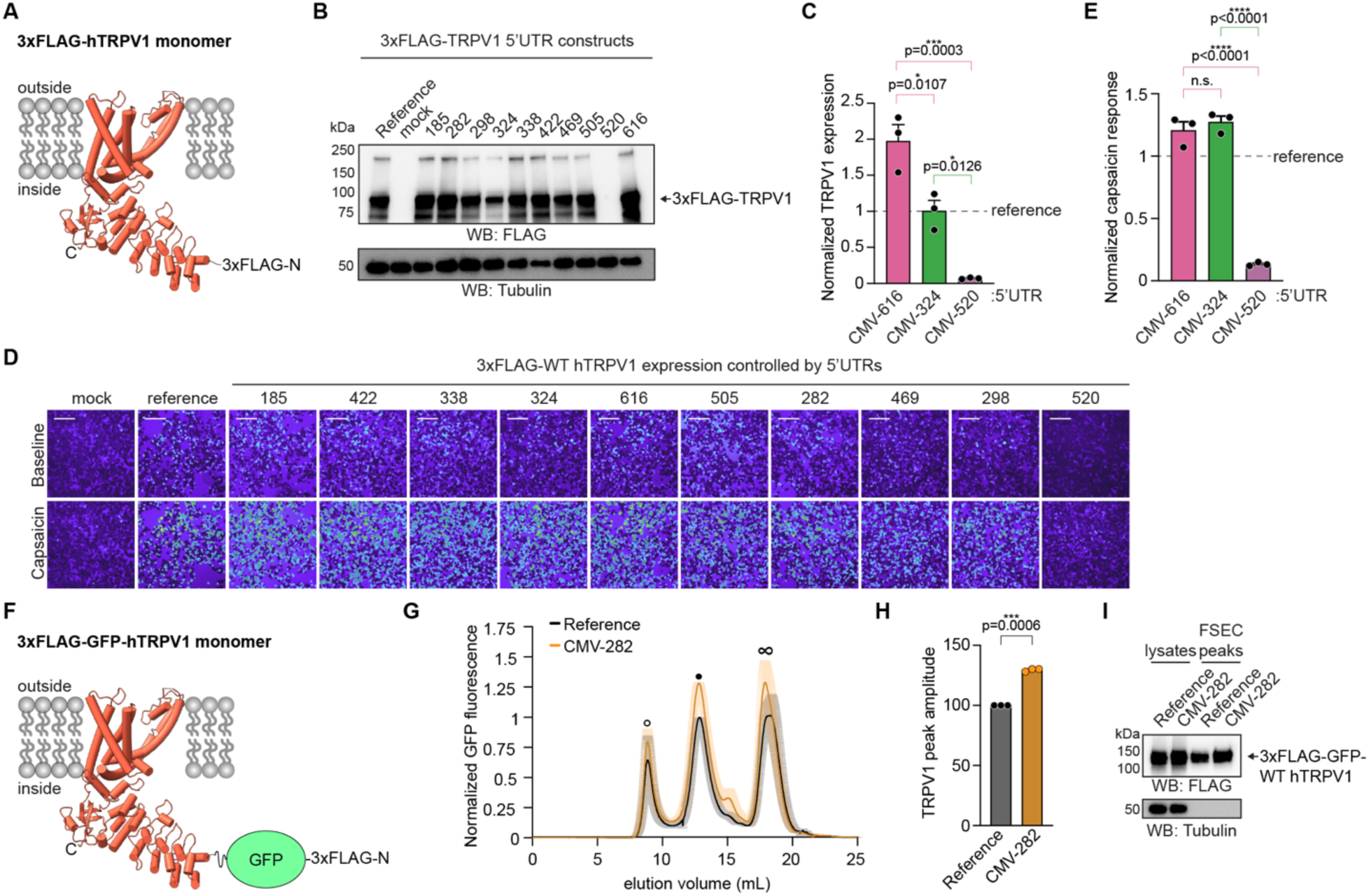
5’UTRs mostly enhance TRPV1 expression and can improve protein production. (**A**) Monomeric ribbon diagram of the human TRPV1 atomic model for residues R115-S781 (PDB: 9P6B); each subunit is 98 kDa. (**B**) Western blot of lysates from cells transfected with empty vector (mock) or expressing 3xFLAG-hTRPV1 controlled by a CMV promoter alone (reference) or with the indicated 5’UTR, probed using HRP-conjugated anti-FLAG antibody. Tubulin was the loading control. Results were verified in three independent experiments. (**C**) Quantification of the reference plasmid-normalized expression of 3xFLAG-hTRPV1 controlled by CMV-616, CMV-324, or CMV-520 from data as in (B). Data represent mean ± SEM. *n* = 3 independent experiments, one-way ANOVA with Tukey’s *post hoc* analysis. (**D**) Ratiometric calcium imaging of HEK293T cells transfected with empty vector (mock), or 3xFLAG-hTRPV1 controlled by CMV alone (reference) or the indicated 5’UTRs. Cells were stimulated with 1 µM capsaicin. Images are representative of three independent experiments and are organized from highest to lowest activity (see Figure S4B). Scale bars indicate 100 µm. (**E**) Quantification of reference plasmid-normalized capsaicin-evoked change in Fura-2 ratio for 3xFLAG-hTRPV1 controlled by CMV-616, CMV-324, and CMV-520. Data represent mean ± SEM. *n* = 3 independent experiments, with an average of *n* ≥ 2,792 cells per transfection condition per biological replicate. P-values listed in figure panel, one-way ANOVA with Tukey’s *post hoc* analysis. (**F**) Monomeric ribbon diagram of the hTRPV1 atomic model from (A) with an N-terminal 3xFLAG-GFP tag diagramed; this construct was used for FSEC analyses. (**G**) FSEC chromatograms from cells expressing 3xFLAG-GFP-hTRPV1 controlled by a CMV promoter only (black) or with CMV-282 (orange). Peaks correspond to void (empty dot), tetrameric hTRPV1 channels (black dot), and free GFP (infinity symbol). Traces normalized to the maximum amplitude of the reference plasmid hTRPV1 peak within each experiment. Data plotted as mean ± SEM. *n* = 3 independent experiments. (**H**) Quantification of the TRPV1 peak amplitude from cells expressing 3xFLAG-GFP-hTRPV1 controlled by the CMV promoter alone (reference) or with CMV-282 from the data in (G). Within each independent experiment, the reference plasmid measurement was set to 100%. *n* = 3 independent experiments, P-value listed in figure panel, two-tailed Student’s t-test. (**I**) Western blot of lysates and TRPV1 peaks (black dot) from samples as in (G) probed using HRP-conjugated anti-FLAG antibody. Tubulin was the loading control. Results were verified in three independent experiments.

Functional measurements using ratiometric Ca^2+^ imaging revealed minimal hTRPV1 activity with CMV-520 and strong activity with all other 5’UTRs and the reference plasmid (**Figures 6D**, **6E**, and **S4B**). CMV-520 exhibited an ∼10-fold reduction in capsaicin-evoked responses, whereas the other constructs produced similar activities, suggesting that hTRPV1 function saturates under these assay conditions (**Figures 6E** and **S4B**).

The ability of most 5’UTRs to elevate hTRPV1 expression above that achieved with the CMV alone is notable, given the widespread use of CMV-driven overexpression for structural studies (Sun et al. 2025; Liao et al. 2013; Cao et al. 2013). Because fluorescently tagged proteins are typically screened by fluorescence size exclusion chromatography (FSEC), we asked whether a 5’UTR could increase hTRPV1 yield in this workflow (Kawate and Gouaux 2006). We generated 3xFLAG-GFP-tagged WT hTRPV1 constructs with or without the CMV-282 5’UTR (**Figures 6F**), transiently expressed them in HEK293T cells, solubilized the proteins in DDM, and analyzed equal amounts of lysate (600 µg) by FSEC. Both constructs produced monodisperse hTRPV1 peaks (**Figures 6G**), but CMV-282 increased peak height by ∼30% across three replicates (**Figure 6H**). Western blotting corroborated higher hTRPV1 production with CMV-282 (**Figure 6I**).

Together, these findings show that 5’UTRs tune hTRPV1 expression across high, intermediate, and low levels, but unlike hTRPA1, the reference plasmid yields an intermediate expression level (**Figure 6C**). Moreover, the CMV-282 5’UTR reproducibly boosts hTRPV1 production from equivalent protein, highlighting its utility for improving yields in structural biology pipelines.

### 5’UTRs mostly enhance TRPM8 expression, revealing distinct effects across open reading frames

Given the disparate effects of the 5’UTRs on hTRPA1 and hTRPV1 expression and function, we next examined how they regulate a third pain receptor, TRPM8 (**Figure 7A**). Like TRPA1 and TRPV1, TRPM8 is a Ca^2+^-permeable non-selective cation channel (Basbaum et al. 2009; Julius 2013). TRPV1 and TRPM8 homotetramers are ∼390 and ∼520 kDa, respectively, making TRPM8 similar in size to hTRPA1; thus, we asked whether protein size contributes to differences in 5’UTR performance.

**Figure 7.**
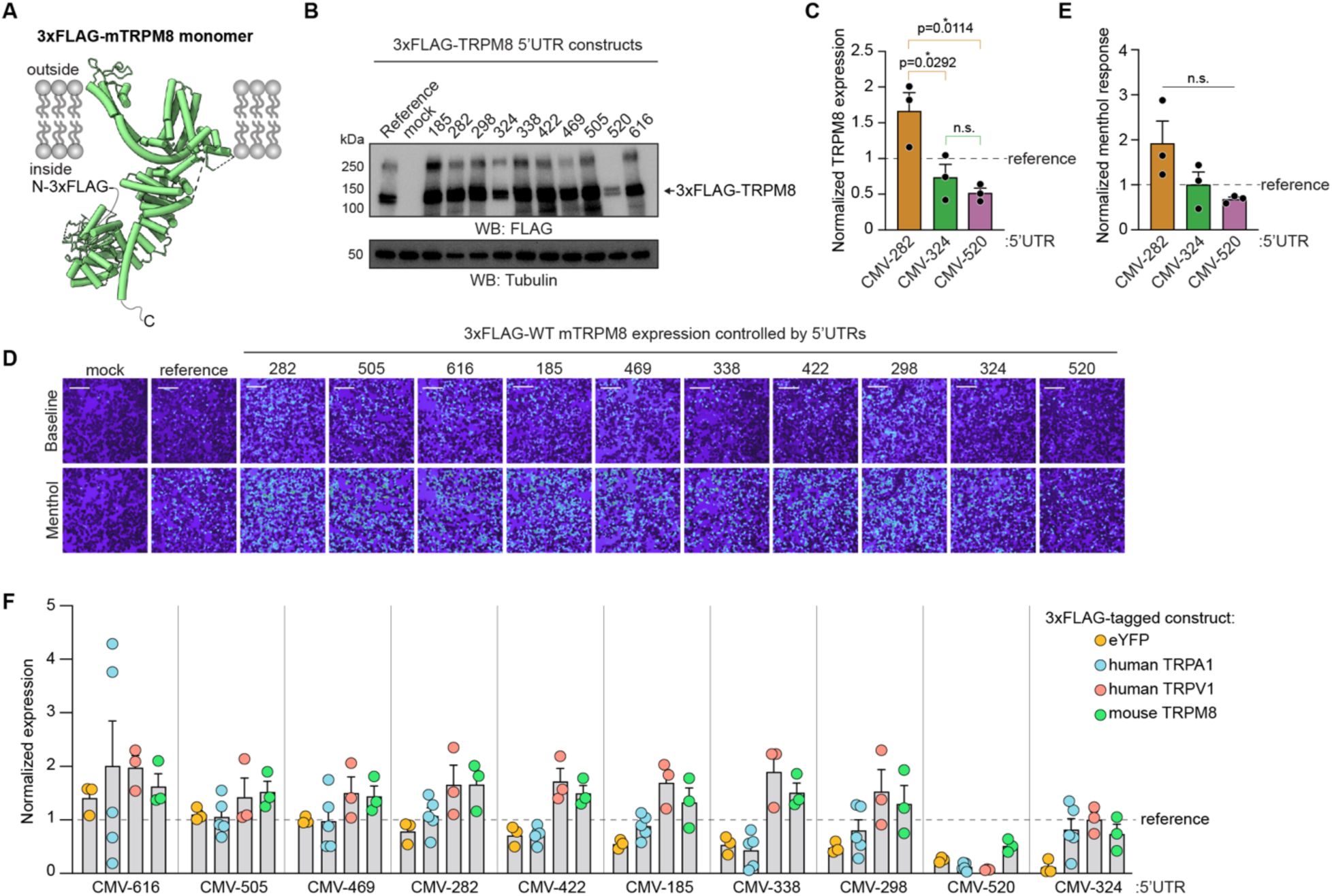
5’UTRs mostly enhance TRPM8 expression, revealing distinct effects across open reading frames. (**A**) Monomeric ribbon diagram of the mouse TRPM8 atomic model for residues D40-K1104 (PDB: 9B6D); each subunit is 130.8 kDa. (**B**) Western blot of lysates from cells transfected with empty vector or expressing 3xFLAG-mTRPM8 controlled by a CMV promoter without (reference) or with the indicated 5’UTR, probed using HRP-conjugated anti-FLAG antibody. Tubulin was the loading control. Results were verified in three independent experiments. (**C**) Quantification of the reference plasmid-normalized expression of 3xFLAG-mTRPM8 controlled by CMV-282, CMV-324, or CMV-520 from data as in (B). Data represent mean ± SEM. *n* = 3 independent experiments, one-way ANOVA with Tukey’s *post hoc* analysis. (**D**) Ratiometric calcium imaging of HEK293T cells transfected with empty vector (mock), or 3xFLAG-mTRPM8 controlled by CMV alone (reference) or the indicated 5’UTRs. Cells were stimulated with 200 µM menthol. Images are representative of three independent experiments and are organized from highest to lowest activity (see Figure S4D). Scale bars indicate 100 µm. (**E**) Quantification of reference plasmid-normalized menthol-evoked change in Fura-2 ratio for 3xFLAG-mTRPM8 controlled by CMV-282, CMV-324, and CMV-520. Data represent mean ± SEM. *n* = 3 independent experiments, with an average of *n* ≥ 2,724 cells per transfection condition per biological replicate. P-values listed in figure panel, one-way ANOVA with Tukey’s *post hoc* analysis. (**F**) Quantification of the reference plasmid-normalized expression of 3xFLAG-tagged eYFP (yellow), hTRPA1 (blue), hTRPV1 (red), or mTRPM8 (green) controlled by the indicated 5’UTR. Data replotted from Figures 3C, 4D, S4A and S4C and arranged from highest-to-lowest for eYFP expression to show differences in 5’UTR control across open reading frames. *n* = 3 (eYFP, hTRPV1, and mTRPM8) or *n* = 5 (hTRPA1) independent replicates.

CMV-520 and CMV-324 decreased or did not alter mouse TRPM8 (mTRPM8) expression relative to the reference plasmid, whereas all remaining 5’UTRs enhanced expression as assessed by anti-FLAG Western blot (**Figures 7B**, **7C**, and **S4C**). Ratiometric Ca^2+^ imaging revealed parallel effects on menthol-evoked activity: CMV-520 slightly reduced channel function, while the remaining 5’UTRs produced stepwise increases (**Figures 7D, 7E**, and **S4D**). Across both expression and functional measurements, the 5’UTRs tuned mTRPM8 expression over a ∼3-fold range between the highest (CMV-282) and lowest (CMV-520) expressors – narrower than the ranges observed for eYFP, hTRPA1, and hTRPV1 (**Figures 7F**). This reduced dynamic range reflects the relatively modest effect of CMV-520 on lowering mTRPM8 levels compared to its stronger effect on hTRPA1 or hTRPV1 (**Figures 7C, D,** and **F**).

Comparison of 5’UTR performance across eYFP, hTRPA1, hTRPV1, and mTRPM8 expression revealed several trends. First, CMV-616 and CMV-520 consistently produced the highest and among the lowest expression levels, respectively (**Figure 7F**). Second, although CMV-324 drove the lowest expression for eYFP, it had only modest effects on the three TRP channels (**Figure 7F**). Third, with few exceptions, eYFP and hTRPA1 displayed a similar range and rank order of 5’UTR-dependent control (**Figure 7F**). Fourth, hTRPV1 and mTRPM8 behaved similarly: all 5’UTRs increased their expression relative to the reference plasmid except CMV-324 and CMV-520 (**Figure 7F**). These patterns highlight distinct 5’UTR effects for eYFP and hTRPA1 versus hTRPV1 and mTRPM8.

Overall, the 5’UTRs provide a versatile toolkit for tuning ectopic protein expression across a broad range. CMV-520 is the most consistent at reducing expression, CMV-616 is the most consistent enhancer, and the remaining 5’UTRs exert gene-specific effects. In the following sections, we illustrate two research contexts in which limiting protein expression with CMV-520 improves signal-to-noise in assays where overexpression can otherwise pose experimental challenges.

### Precision-ID: Controlled TurboID expression significantly improves labeling specificity

TurboID is a proximity biotinylation tag widely used to identify protein interactions in cells (Cho et al. 2020). Our lab recently showed that calmodulin (CaM) functions as a hTRPA1 auxiliary subunit that pre-associates with the channel at rest to tightly regulate its activity during Ca^2+^ influx (Sanders et al. 2025; Quevedo et al. 2025). To demonstrate this complex in cells, we previously fused TurboID to the CaM C-lobe and compared biotinylation of WT versus CaM-binding deficient hTRPA1 channels (**Figure 8A**) (Sanders et al. 2025). Although these experiments revealed greater WT than mutant biotinylation, the mutants were still strongly biotinylated – likely due to TurboID overexpression – which limited the specificity of the assay and reduced confidence in the conclusions.

**Figure 8.**
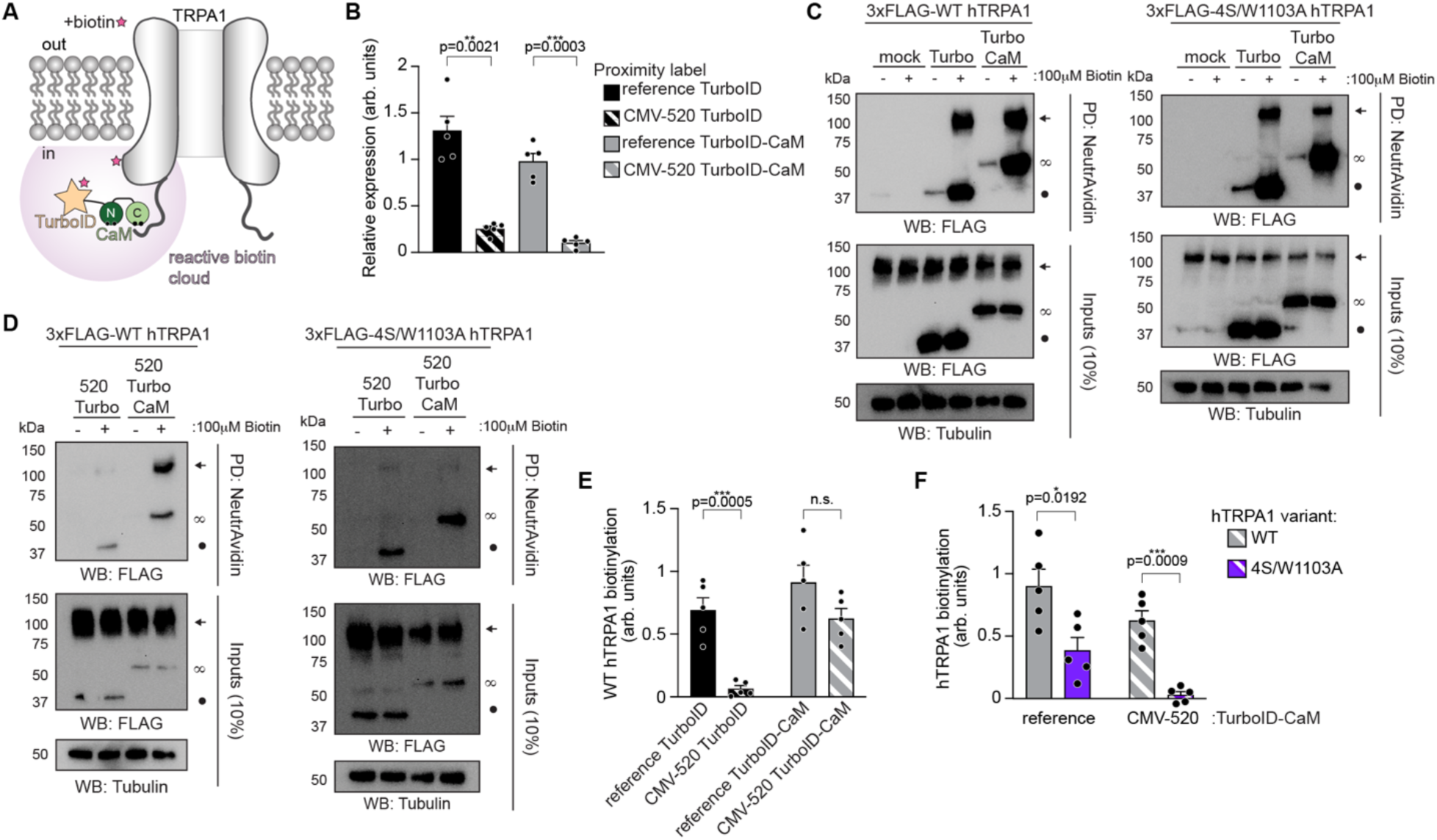
Controlled TurboID expression improves labeling specificity: Precision-ID. (**A**) Approach for proximity biotinylation. TurboID (orange star) fused to CaM (green) is co-expressed with hTRPA1 variants (gray) in cells. Addition of biotin (pink star) to the media will facilitate generation of a local reactive cloud (purple) that will biotinylate TurboID-CaM and TRPA1, pending an interaction. (**B**) Quantification of the relative expression of 3xFLAG-tagged TurboID free tag (black) or 3xFLAG-TurboID-CaM (grey) controlled by a CMV promoter without (reference, solid bars) or with CMV-520 (striped bars) from data as in (C and D). *n* = 5 independent experiments. (**C** and **D**) Immunoblotting analysis of biotinylated 3xFLAG-WT or 4S/W1103A hTRPA1 co-expressed with empty vector (mock), or 3xFLAG-TurboID or 3xFLAG-TurboID-CaM controlled by (C) a CMV promoter alone (reference) or (D) with CMV-520 in HEK293T cells. Neutravidin immunoprecipitated eluates and whole cell lysate inputs were probed with anti-FLAG antibody. Tubulin was the loading control. Blots are representative of five independent experiments. (**E**) Quantification of the relative biotinylation (arb. units) of WT hTRPA1 by TurboID controlled by a CMV promoter only (solid black) or with CMV-520 (black stripe), or by TurboID-CaM controlled by a CMV promoter only (solid grey) or with CMV-520 (grey stripe) from data as in (C and D). *n* = 5 independent experiments. (**F**) Quantification of the relative biotinylation of WT (grey) or 4S/W1103A (purple) hTRPA1 by TurboID-CaM controlled by a CMV promoter only (reference, solid bars) or with CMV-520 (striped bars) from data as in (C and D). *n* = 5 independent experiments. (B, E, and F) Data represent mean ± SEM. P-values listed in figure panel. Two-tailed Student’s t-tests.

Therefore, we asked whether CMV-520 could improve labeling specificity by lowering TurboID expression. We generated 3xFLAG-tagged free TurboID and TurboID-CaM constructs with and without CMV-520, and anti-FLAG Western blot confirmed reduced construct expression with the 5’UTR (**Figures 8B-D**). We then examined how CMV-520 influenced TurboID-CaM biotinylation of WT hTRPA1 versus a CaM-binding deficient mutant (4S/W1103A) (Quevedo et al. 2025). Without the 5’UTR, both free TurboID and TurboID-CaM strongly biotinylated WT and 4S/W1103A channels (**Figure 8C**, arrows). As in our prior work, TurboID-CaM selectively biotinylated WT channels more than free TurboID (**Figure 8E**, compare solid bars) and biotinylated the 4S/W1103A mutant less efficiently (**Figure 8F**, solid bars).

In contrast, inclusion of CMV-520 markedly sharpened labeling specificity. TurboID-CaM but not free TurboID strongly biotinylated WT hTRPA1 (**Figure 8D** and **E**, compare striped bars), and TurboID-CaM produced almost no biotinylation of the 4S/W1103A mutant (**Figure 8D** and **F**, striped bars).

Together, these results show that restricting TurboID expression with CMV-520 substantially improves WT hTRPA1-specific biotinylation by TurboID-CaM while eliminating off-target labeling of CaM-binding deficient channels (**Figure 8F**, striped versus solid bars). These findings not only strengthen our conclusion that CaM pre-associates with WT hTRPA1 in resting cells (Sanders et al. 2025), but they circumvent the main issue of unspecific proximity interactions with overexpressed TurboID-like tools. The increased selectivity afforded by controlling biotin-conjugating proximity tag expression with CMV-520, which we term “Precision-ID”, will be applicable for interaction studies across all biological fields.

### Reduced G3BP1 expression improves its use as a stress granule marker

The Ras GTPase-activating protein-binding protein 1 (G3BP1) is an RNA-binding protein important for the assembly and dynamics of stress granules (SG), cytosolic protein-RNA selective phase environments that form in response to cellular stress (Omer et al. 2020; Freibaum et al. 2024). G3BP1 is widely used as an SG biomarker readout in drug screens aimed at identifying therapeutic compounds for frontotemporal dementia (FTD) and amyotrophic lateral sclerosis (ALS), among other neurodegenerative disorders (Freibaum et al. 2024; Mahajan et al. 2025; Fang et al. 2019; Uechi et al. 2025). However, G3BP1 readily forms cytoplasmic aggregates when overexpressed, even in the absence of stress, which complicates experiments or screens that require long expression times (Guillén-Boixet et al. 2020). Moreover, because aggregates formed by protein overexpression do not necessarily reflect the disease-relevant SG state, many viable small-molecule candidates may be overlooked. Such compounds may fail to reverse cytosolic aggregates driven by overexpression artifacts yet remain fully capable of modulating SGs under more physiological conditions. Although inducible stable cell lines can temporally control G3BP1 levels, they are sensitive to identifying false-positives that inadvertently inhibit protein production and fail to solve overexpression issues in assays requiring extended timepoints.

Therefore, we asked whether CMV-520 could reduce G3BP1 expression during transient transfection to prevent overexpression-driven aggregate formation and thereby improve its performance as a stress-dependent SG marker. We generated GFP-tagged G3BP1 constructs with and without CMV-520 (**Figure 9A**). Transient expression in HEK293T cells for 24 hours revealed that CMV-520 significantly lowered G3BP1 abundance as measured by anti-GFP Western blotting (**Figure 9B**). To assess whether reduced expression mitigated aggregation, we transiently transfected HeLa cells with GFP-G3BP1 with and without CMV-520, fixed them after 24 hours, and quantified the number of GFP-G3BP1 granules by epifluorescence microscopy (**Figure 9C**). Cells expressing GFP-G3BP1 under CMV-520 showed markedly fewer cytoplasmic GFP-G3BP1 granules than those expressing the reference construct (**Figures 9C** and **D**, without stressor), indicating that the 5’UTR alleviates overexpression-induced ectopic G3PB1 accumulation.

**Figure 9.**
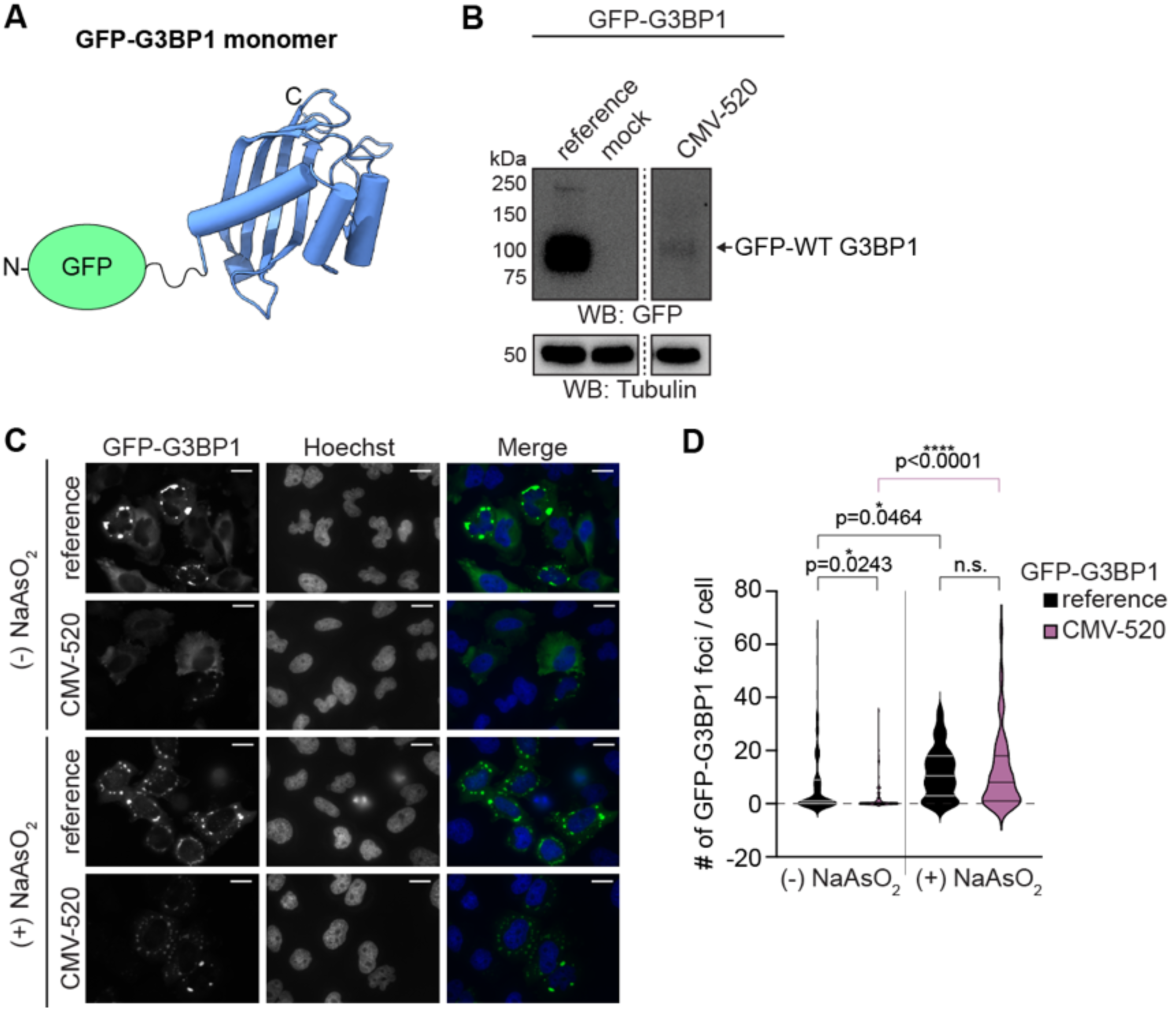
Reduced G3PB1 expression improves its utility as a stress granule marker. (**A**) Monomeric ribbon diagram of G3BP1 atomic model (PDB:8VL1) with an N-terminal GFP tag diagramed. Untagged G3BP1 produces a 54.6 kDa protein. (**B**) Western blot of lysates from cells transfected with empty vector (mock) or expressing GFP-G3BP1 controlled by a CMV promoter only (reference) or with CMV-520 probed using an anti-GFP antibody. Tubulin was the loading control. (**C**) Representative epifluorescence images of HeLa cells expressing GFP-G3BP1 controlled by a CMV promoter only (reference) or with CMV-520 treated with or without 0.5 mM sodium arsenite (NaAsO_2_). Cells were stained with Hoechst (blue). Scale bar indicates 4 µm. (**D**) Violin plot quantification of GFP-G3BP1 foci (granules) per cell for data as seen in (C). Each condition was tested with an *n* ≥ 80 cells per transfection condition. P-values listed in figure panel, one-way ANOVA with Tukey’s *post hoc* analysis.

We next tested whether reduced expression improved the utility of GFP-G3BP1 as an SG marker by treating cells with sodium arsenite (NaAsO_2_), a commonly used reagent to induce SG formation (Sahoo et al. 2018). Strikingly, despite the significantly decreased background observed in the CMV-520 condition, treatment with NaAsO_2_ resulted in the same number of GFP-G3BP1 granules in both the CMV-520 and reference plasmid constructs (**Figures 9C** and **D**, with stressor). Because CMV-520 markedly reduced baseline granules while preserving stress-induced granule formation, it substantially improved the signal-to-noise ratio for SG detection (**Figures 9C** and **D**, purple plots).

Together, these results show that lowering G3BP1 expression with CMV-520 enhances the specificity and performance of GFP-G3BP1 as a stress-induced SG biomarker in transiently transfected cells, which will be valuable for experiments and drug screen campaigns requiring long transfection times.

## DISCUSSION

Our study establishes a modular and scalable strategy to tune protein abundance during transient transfection using short (<170 bp) 5’UTRs. By integrating a panel of previously identified 5’UTRs (Lewis et al. 2025) into constructs driven by a CMV promoter, we defined a reproducible dynamic range of protein expression across fluorescent reporters and large membrane proteins. We benchmarked these 5’UTRs against five commonly used eukaryotic promoters and showed that for eYFP and hTRPA1, 5’UTRs provide finer, more continuous control over expression levels than promoter swapping. Finally, we demonstrated the utility of this system across diverse experimental contexts – including TurboID-based proximity labeling, ion channel functional assays, protein production, and stress granule biology – highlighting how tunable expression enables clearer, more physiologically relevant readouts.

Precise control over protein expression is essential because different experimental goals demand different abundance thresholds (Mori et al. 2020; Moriya 2015). High expression can be advantageous for applications such as protein purification, structural studies, or difficult-to-detect proteins. Intermediate expression may be necessary for assays that require robust signal without saturating downstream pathways. In contrast, low expression is critical for maintaining near-endogenous stoichiometry in trafficking studies, protein-protein interaction mapping, or any system where overexpression induces artifacts, cytotoxicity, or altered localization. The ability to select a 5’UTR that reliably positions a protein within these expression windows provides a powerful alternative to trial-and-error optimization of DNA dose, transfection duration, or promoter strength (Qin et al. 2010; Zhao et al. 2020).

We also observed protein-specific differences in how 5’UTRs modulated expression, particularly among the three TRP ion channels tested. hTRPA1, hTRPV1, and mTRPM8 displayed ∼10-fold, ∼25-fold, and ∼3-fold expression ranges, respectively. Additionally, while the 5’UTRs tuned hTRPA1 abundance across a broad gradient, most 5’UTRs enhanced hTRPV1 and mTRPM8 expression compared to a CMV promoter alone. Together, these findings reveal that 5’UTRs do not function as universal “boosters” or “dampeners” but integrate protein coding sequences and mRNA folding landscapes to influence translation efficiency. Such context dependency opens opportunities for future work to dissect how 5’UTR motifs, codon usage, RNA secondary structure, and ribosome engagement collectively shape expression – offering a route to new discoveries in RNA biology and synthetic control of translation.

Functionally, we show that these 5’UTRs enable clearer interpretation of assays prone to artifacts from overexpression. The lowest-expressing 5’UTR across all proteins tested (CMV-520) enhanced the detection of functional differences in a hyperactive hTRPA1 channel mutant (**Figure 5**), substantially suppressed background labeling in TurboID proximity biotinylation assays yielding an improved strategy that we call “Precision-ID” that will be broadly applicable to future interaction mapping applications across biological fields (**Figure 8**), and reduced aggregation of overexpressed G3BP1 while preserving drug-induced stress granule formation thereby increasing the specificity of this biomarker for future drug screen campaigns (**Figure 9**). Conversely, a 5’UTR that enhanced hTRPV1 expression (CMV-282) improved protein yields in FSEC analyses (**Figure 6G-I**), demonstrating that 5’UTRs can also be leveraged to boost expression when higher protein abundance is advantageous.

Looking forward, the 5’UTR toolkit presented here offers a simple, compact, and highly modular method to fine-tune protein expression in transient transfection systems. Because these elements are short and easy to clone, they can be rapidly incorporated into diverse constructs, applied across proteins of different sizes and membrane complexities, and adapted to a wide range of mammalian cell types and experimental platforms. As molecular biology continues to push toward more physiologically grounded experimental systems, scalable tuning of protein expression will be increasingly valuable. These 5’UTRs thus provide a practical and versatile framework for improving experimental precision, enabling new mechanistic discoveries, and facilitating the design of next-generation RNA-based regulatory tools.

### Limitations of the study

While the 5’UTR panel provides a practical and modular strategy to tune protein abundance, there are limitations that should be considered. Across four diverse proteins – from cytosolic reporters to large ion channels – we found that only CMV-616 and CMV-520 consistently yielded among the highest and lowest expression levels, respectively. The eight remaining 5’UTRs showed protein-specific effects, suggesting that translation efficiency is influenced by sequence-dependent interactions between each 5’UTR and the downstream open reading frame. Consequently, researchers may need to empirically test the full 5’UTR panel for any new protein target to identify the most suitable expression level for their applications. In practice, this screening process was straightforward due to our use of a common MegaPrimer-based cloning workflow and the ability to easily swap open reading frames (**Figure S2**).

Eukaryotic promoters, by comparison, produced predictable high or low expression independent of protein identity but offered only coarse control and required more labor-intensive cloning (Forrest et al. 2014; Qin et al. 2010). Importantly, promoter choice and 5’UTR selection may not be mutually exclusive. Future applications may combine specific 5’UTRs with alternative eukaryotic promoters to achieve even finer gradations in protein expression, enabling more precise tuning of experimental conditions.

## MATERIALS AND METHODS

### Plasmid Construction

Cloning for 3xFLAG-tagged human TRPA1, human TRPV1, mouse TRPM8, and 4S/W1103A human TRPA1 in CMV containing p3xFLAG-eYFP vector was reported previously (Bali et al. 2023) (Quevedo et al. 2025). 3xFLAG-eYFP/hTRPA1 controlled by the EF1α, PGK, UbC, or SV40 promoters were created by first introducing two XbaI sites flanking the CMV enhancer and promoter by site-directed mutagenesis (Agilent). Each promoter was then introduced by InFusion cloning (Takara) into XbaI-digested 3xFLAG-eYFP/hTRPA1. The 5’UTRs were PCR amplified from plasmids generously provided by Wendy Gilbert and Carson Thoreen using a common CMV enhancer forward primer and a 5’UTR-specific reverse primer that contained the beginning of the 3xFLAG tag (**Figure S2**). The resulting MegaPrimers were gel purified and used to introduce each 5’UTR into the 3xFLAG vector through site-directed mutagenesis. Production of 3xFLAG-TurboID was reported previously (Sanders et al. 2025). 3xFLAG-TurboID-CaM was built from a 3xFLAG-TurboID-CaM C-lobe construct reported previously (Sanders et al. 2025) by site-directed mutagenesis with a MegaPrimer including part of TurboID, the CaM N-lobe, and part of the CaM C-lobe. 3xFLAG-GFP-hTRPV1 was built by site-directed mutagenesis with a MegaPrimer including part of CMV enhancer + promoter and the 3xFLAG tag. CMV-520 GFP-G3BP1 was built using site-directed mutagenesis to add the CMV-520 sequence.

All DNA primers were ordered from ThermoFisher and all constructs were sequence verified using the Yale School of Medicine Keck DNA Sequencing Facility. Additional cloning details including primers used and the sequences for the eukaryotic promoters and 5’UTRs are provided in the accompanying Source Data file.

### Mammalian Cell Culture and Protein Expression

Human embryonic kidney cells (HEK293T, ATCC CRL-3216) were cultured in Dulbecco’s modified Eagle’s medium (DMEM; Invitrogen) supplemented with 10% calf serum and 1x Penicillin Streptomycin (Invitrogen) at 37 °C and 5% CO_2_. Cells were grown to ∼85-95% confluence before splitting for experiments or propagation. HEK293T cells cultured to ∼95% confluence were seeded at 1:10 or 1:20 dilutions into 6- or 12-well plates (Corning), respectively. After 1-5 hours recovery, cells were transiently transfected with 1-2 μg plasmid using jetPRIME (Polyplus) according to manufacturer protocols.

Neuro-2A cells (N2A, ATCC CCL-131) were cultured in Dulbecco’s modified Eagle’s medium (DMEM + GlutaMAX; Invitrogen) supplemented with 10% fetal bovine serum, 1x non-essential amino acids and 1x Penicillin Streptomycin (Invitrogen) at 37 °C and 5% CO_2_. Cells were grown to ∼85-95% confluence before splitting for experiments or propagation. N2A cells cultured to ∼95% confluence were seeded at 1:10 or 1:20 dilutions into 6- or 12-well plates (Corning), respectively. After 2-5 hours recovery, cells were transiently transfected with 1-2 μg plasmid using Lypofectamine2000 (Invitrogen) according to manufacturer protocols.

Human epithelial cells (HeLa, ATCC CCL-2) were cultured in Dulbecco’s modified Eagle’s medium (DMEM; Invitrogen) supplemented with 10% fetal bovine serum and 1x Penicillin Streptomycin (Invitrogen) at 37 °C and 5% CO_2_. Cells were grown to ∼85-95% confluence before splitting for experiments or propagation. HeLa cells cultured to ∼95% confluence were seeded at 1:10 into a 24-well plate (Corning). After 2-5 hours recovery, cells were transiently transfected with 1-2 μg plasmid using Lypofectamine3000 (Thermo Fisher Scientific) according to manufacturer protocols.

### Ratiometric Ca^2+^ Imaging

40-48 hours post-transfection, HEK293T cells were plated into isolated silicone wells (Sigma) on poly-L-lysine (Sigma)-coated cover glass (ThermoFisher). Remaining cells were lysed for anti-FLAG immunoblotting to ensure equivalent expression, as detailed below. After adhering to the glass slide for 1 hour, cells were loaded with 10 μg/mL Fura 2-AM (ION Biosciences) in physiological Ringer’s solution (in mM: 120 NaCl, 5 KCl, 2 CaCl_2_, 25 NaHCO_3_, 1 MgCl_2_, 5.5 HEPES, 1 D-glucose, pH 7.4; Boston BioProducts) with 0.025% Pluronic F-127 (Sigma) and 1% DMSO, and incubated for 45 minutes at 37 °C, then rinsed twice with Ringer’s solution. Ratiometric Ca^2+^ imaging was performed using a Zeiss Axio Observer 7 inverted microscope with a Hamamatsu Flash sCMOS camera at 20x objective. Dual images (340 and 380 nm excitation, 510 nm emission) were collected and pseudocolour ratiometric images were monitored during the experiment (MetaFluor software). After stimulation with agonist, cells were observed for 50 seconds. AITC, capsaicin, and menthol were purchased from Sigma and were freshly prepared as a stock at 4x the desired concentration in 1% DMSO and Ringer’s solution. 5 μL of 4x agonist was added to wells containing 15 μL Ringer’s solution to give the final 1x desired concentration. For all experiments, a minimum of 60-90 cells were selected per condition per replicate for ratiometric fluorescence quantification in MetaFluor with 5 replicates per experiment. Background signal was quantified from unresponsive cells and subtracted from quantified cells for normalization.

### Cell Lysis and Pulldown Experiments

#### Whole cell lysates – calcium imaging experiments

Resuspended HEK293T cells were pelleted at 1000 rpm for 5 min at room temperature. Media was removed, the cells were washed once with 1x PBS (Ca^2+^ and magnesium free, Boston Bioproducts) and lysed in 100 μL of lysis buffer (40 mM Tris pH 8.0, 150 mM NaCl, 5 mM DDM, cOmplete protease cocktail inhibitor tablet) at 4 °C while gently nutating. Cell debris were pelleted from the resulting lysates by centrifugation at 15,000 RPM for 10 minutes at 4 °C. Total protein concentration in lysates were quantified using a BCA assay (Pierce). Equal concentrations of protein lysate (100 μg) from each experimental condition were analyzed.

#### Biotinylation Assays

For proximity biotinylation assays, HEK293T cells co-expressing 3xFLAG-TurboID or 3xFLAG-TurboID-CaM and 3x-FLAG-tagged WT or 4S/W1103A hTRPA1. After ∼20 hours, cells were treated with 100 μM biotin for 10 minutes at 37 °C to induce proximity biotinylation. Media was then removed, cells were washed once with 1x PBS, and cells were lysed as described above. Equal concentrations of protein lysate (100 µg) were affinity purified via protein biotinylation as described below.

#### NeutrAvidin pulldowns

Cell lysates that were generated following biotin labeling. Lysates were incubated with 15 μL of lysis buffer-equilibrated Neutravidin resin (Pierce) for 1 hr at 4 °C with gentle nutation. The resin was then washed with lysis buffer three times, followed by two harsher washes with 1× PBS + 100 mM DTT. Biotinylated protein was eluted from the resin with a multi-step protocol to prevent TRPA1 aggregation while maximizing protein elution from the resin. First, resin was incubated with 10 μL of biotin elution buffer (TRPA1 lysis buffer, 100 mM Glycine, 10 mM Biotin, 1% SDS) on ice for 10 min, followed by addition of 1 μL ß-mercaptoethanol (BME; Sigma) to each sample and incubation on ice for 5 min, and finally by addition of 4 μL 6× Laemmli buffer + 10% BME with incubation at 65 °C for 10 min. The resin was centrifuged; supernatant was removed and combined with additional 4 μL Laemmli buffer + 10% BME for SDS-PAGE analysis.

### SDS-PAGE and Immunoblot

Samples were combined with 6x reducing loading dye (Boston Bioproducts) and separated on precast 4-20% SDS-PAGE gels (Bio-Rad). Gels were transferred onto PVDF membranes (Bio-Rad) by semi-dry transfer at 15V for 30 minutes. Blots were blocked in 3% BSA or 5% milk prior to antibody probing. The following primary antibodies were used in 1x PBST buffer (Boston Bioproducts): anti-FLAG (mouse, 1:30,000, Sigma), anti-GFP (mouse, 1:5,000 in 5% milk, Takara), and anti-tubulin (mouse, 1:5,000 in 3% BSA, Sigma). HRP-conjugated IgG secondary anti-mouse antibody was used as needed (rabbit, 1:50,000, Invitrogen). Membranes were developed using Clarity Western ECL substrate (Bio-Rad) and imaged using a ChemiDoc Imaging System (Bio-Rad). Densitometric quantifications were performed with ImageJ software. All quantified band intensities for eluted samples were divided by their tubulin-normalized input band intensities. All uncropped blots are presented in Figures S5 and S6. Antibody dilutions used were as previously reported (Quevedo et al. 2025) and specificity was further established by 1) clean band signal only or primarily at expected sizes in the imaged blots (Figures S5 and S6) and 2) no signal in expected negative control lanes in pulldown experiments (*e.g.*, Figure 8C-D).

### Fluorescence Size Exclusion Chromatography (FSEC)

FSEC procedures were adapted from those previously reported (Bali et al. 2023; Kawate and Gouaux 2006). Briefly, sample volumes were brought to 600 µL with lysis buffer. Total protein concentration in lysates was quantified using a BCA assay (Pierce). Equal concentrations of protein lysate (600 μg) from each experimental condition were passed through 0.22-micron filters (Costar) by centrifugation, loaded onto a Superose 6 Increase column (GE Healthcare) pre-equilibrated with FSEC buffer (50 mM HEPES, 150 mM NaCl, 1 mM DTT, 0.5 mM DDM, pH 8), and run at a flow rate of 0.5 ml/min. The in-line fluorescence detector (Shimadzu) spectral settings were as follows: Ex:488; Em: 510 nm (eGFP); time increment, 1 s; integration time, 1 s. Data were collected with Unicorn v7.2 software. Fractions corresponding to TRPV1 peaks were collected, concentrated using Amicon Ultra 100 kDa cut-off spin filters, and subjected to immunoblot analysis. For experiments in Figures 6G-I, GFP-tagged CMV-282 and reference plasmid constructs were expressed for 40-48 hours.

### Immunofluorescence staining and imaging

Neuro2A cells (ATCC CCL-131) were transiently transfected as outlined above with 3xFLAG-hTRPA1 plasmids containing the suite of eukaryotic promoters or 5’UTRs and a plasmid encoding soluble mCherry under a CMV promoter. These cells were transferred to poly-L-lysine-coated cover slips and incubated for 16–20 h prior to immunostaining. Cells were fixed on coverslips with 4% paraformaldehyde for 10 min, then permeabilized with 0.1% TritonX-100 in PBS for 10 min. Cells were then blocked in 5% BSA + 2% Glycine in PBS for 5 minutes followed by incubation in primary anti-TRPA1/ANKTM1 (mouse, 1:100, Santa Cruz biotechnologies sc-376495) in 5% BSA + 2% Glycine in PBS at room temperature for 1 hr. Cells were then washed with PBS and incubated with Hoechst 33342 (1:2000, Invitrogen) in 5% BSA + 2% Glycine in PBS at room temperature for 5 minutes. Representative images in Figures 2 and Figure 4 were acquired on a Zeiss Axio Observer Z1 inverted microscope equipped with an Airyscan detection unit, using a Plan-Apochromat 63x/1.40 NA oil objective.

#### High-content Imaging

Widefield images were acquired using the ImageXpress 4.0 high-throughput fluorescent imaging system (Molecular Devices) at the Yale Center for Molecular Discovery. 36 fields of view per well were acquired at 20X magnification with 2 × 2 pixel binning using DAPI, GFP, and Texas Red channels to visualize HOECHST (nucleus), GFP, and mCherry signals, respectively.

#### Image analysis

Images were analyzed using Custom Module Editor of the MetaXpress high content image analysis software (Molecular Devices, Version 6). Briefly, nuclei were segmented based on the HOECHST staining. Cytoplasm was defined as a 2-pixel perinuclear ring, and average nuclear and cytoplasmic mCherry and GFP intensities were quantified in individual cells. Histograms showing the distribution frequency of nuclear or cytoplasmic mCherry average intensity values across individual cells were plotted and used to select a threshold value to define mCherry-positive cells.

Histograms showing the distribution frequency of the nuclear and cytoplasmic average GFP intensity values and normalized GFP/mCherry intensity ratios in the transfected cells were then plotted for all cells positive for mCherry expression.

#### G3BP1-GFP stress granule quantification

HeLa cells were transfected with either G3BP1-GFP (Addgene #135997) or CMV-520-G3BP1-GFP using. After 24 h, cells were treated with or without NaAsO_2_ (0.5 mM) and incubated for 30 min. Cells were washed with PBS, fixed with 4% paraformaldehyde (PFA) for 20 minutes, permeabilized with 0.1% Triton X-100 for 5 minutes, and stained with Hoechst 33342 dye (1:2000, Invitrogen, H1399). Images were captured using a Zeiss AXIO Observer Z1 microscope with a ×63/1.40 oil objective.

To quantify the number of G3BP1-GFP granules, we used an analogous method previously employed to detect nuclear condensates (Poch et al. 2025). Briefly, two CellProfiler pipelines were constructed with slight differences in the minimum threshold values for the CMV-520 G3BP1-GFP variant compared with reference plasmid G3BP1-GFP. This was necessary due to significant differences in total fluorescence intensity between each condition. Both pipelines are available at: https://github.com/dylanpoch/ca-quant.git. The nuclei of each cell were detected before quantifying the overall G3BP1 intensity to filter out untransfected cells. The small G3BP1-GFP granules were identified (with background suppression) and merged with any large G3BP1-GFP granules (diameter > 30 pixels, without background suppression) using three-class Otsu adaptive thresholding. Measurements for the total number and size/intensity of each granule were recorded and exported to R. After grouping cells into their respective conditions, the number of G3BP1-GFP granules per condition was compared using a one-way ANOVA with Tukey’s *post hoc* analysis.

### Plate Reader Assay

16-24 hours post-transfection, HEK293T cells were resuspended in 2 mL of media. For each construct, live-cell count was determines using TC-20 Automated cell counter (Bio-Rad). To a 96-well plate treated with poly-L-Lysine (Sigma), 500,000 cells were added into 100 µL of media. Cells were incubated for 1 hr at 37°C. eYFP fluorescence was measured with a SYNERGYMx plate reader (BioTek) with the fluorescence detector spectral settings set to Ex:485; Em: 528 nm (eYFP) with a gain of 50.

### Automated analysis pipeline for quantifying calcium imaging data

We developed a custom analysis pipeline to automate ratiometric calcium imaging at both the single-cell and population levels. Pseudocolored ratiometric images were acquired using the MetaFluor software and exported as .TIF files at both 5 and 45 second timepoints. Using CellProfiler 4.2.8, an open-source cellular image quantification software (Carpenter et al. 2006), we customized a pipeline to extract the green and blue channels from multicolored MetaFluor images into separate, grayscale images. To ensure all cells were being identified despite temporal changes in fluorescence intensity, single cells were identified from both the green and inverted blue channels using three-class Otsu adaptive thresholding with a diameter of between 25-150 pixels, before merging any overlapping cells identified by both images. The fluorescence intensity (green channel) indicative of intracellular calcium was quantified before exporting the data in .csv files. For details on the exact parameters used, please see our open-source code available at: https://github.com/dylanpoch/ca-quant.git.

Data processing was performed in *R Studio* (Version 2023.12.1+402) using the tidyverse ecosystem (Wickham et al. 2019) including dplyr for data manipulation, tidyr for reshaping, and stringr for parsing filenames. After grouping cells by replicate and condition, the ratio of expressing cells was calculated by taking the number of cells at or above the fluorescence threshold limit (identified with CellProfiler) divided by the total number of cells. All ratios were calculated using images taken from the representative 45 second timepoint. Optional readouts include the mean fluorescence intensity and the integrated intensity per cell, among others.

### Statistical Analysis

All data quantification was performed in Microsoft Excel. Quantified data presentation and statistical analyses were performed in GraphPad Prism. In most experiments (indicated by “normalized” in figure legends and figure panels), data from constructs controlled by a CMV promoter without a 5’UTR (“reference plasmids”) were used to normalize all other samples within individual biological replicates. In those quantified data, the dashed line at 1 indicates the reference plasmid value. Criterion for statistical significance for all tests was p<0.05. The statistical tests applied are indicated in figure legends. P-values are listed in figure panels.

## DATA AVAILABILITY

The data that support the findings of this study are available as an accompanying Source Data file. Uncropped blots are provided in Figures S5 and S6 in the accompanying Supplemental Material document.

The 3V3D (https://www.rcsb.org/structure/3V3D), 6PQQ (https://www.rcsb.org/structure/6PQQ), 9P6B (https://www.rcsb.org/structure/9P6B), 9B6D (https://www.rcsb.org/structure/9B6D), and 8V1L (https://www.rcsb.org/structure/8V1L) PDB files were used in this study. Models were built with ChimeraX version 1.10.1.

## ACKNOWELDGEMENTS

We thank Wendy Gilbert, Tony Koleske, Steven Tang, and Christian Schlieker for constructive suggestions, members of the Paulsen lab for helpful discussions, and Grover Paulsen-Sharpe and Marilee Pawsen for moral support. We thank the Yale Center for Molecular Discovery for their assistance with high-content imaging and image data analysis. The core is supported in part by an NCI Cancer Center Support Grant #NIH P30 CA016359. This work utilized ImageXpress Micro 4 high content imager that was purchased with funding from a National Institutes of Health SIG grant 1S10OD032384. We thank the Yale Imaging and Flow Cytometry Facilities on Science Hill.

C.G. is supported by a Biophysics pre-doctoral training grant (T32GM149438) and Gruber Science Fellowship from Yale. D.P. is supported by a Cellular, Molecular, and Quantitative Biology Training Program pre-doctoral training grant (). A.M.A. is supported by a Chemical Biology pre-doctoral training grant (T32GM149444). This research was supported by NIH grant R35GM142825 to C.E.P. The content is solely the responsibility of the authors and does not necessarily represent the official views of the National Institutes of Health.

## AUTHOR CONTRIBUTIONS

C.G. and C.E.P. planned the project, designed experiments and cloned most constructs used in the study. A.M.A. cloned and validated the TurboID tools. C.G. conducted all experiments except D.P. performed the G3BP1 HeLa cell work and epifluorescence imaging. D.P. generated and applied tools to quantify G3BP1 aggregation and ratiometric calcium imaging. C.G. and C.E.P. wrote the manuscript with input from D.P. and A.M.A.

## Abbreviations

5’UTR: 5’ untranslated region
CMV promoter: cytomegalovirus promoter
EF1α promoter: elongation factor 1-alpha promoter
SV40 promoter: simian virus 40 promoter
PGK promoter: phosphoglycerate kinase 1 promoter
UbC promoter: ubiquitin C promoter
Ca^2+^: Calcium
eYFP: enhanced yellow fluorescent protein
TRPA1: Transient receptor potential ankyrin 1
TRPV1: Transient receptor potential vanilloid 1
TRPM8: Transient receptor potential melastatin 8
G3BP1: Ras GTPase-activating protein-binding protein 1

