## Supplemental Figures for "5’Untranslated regions provide a versatile toolkit for tunable exogenous protein expression"

•Supplemental Material Figures S1-S6

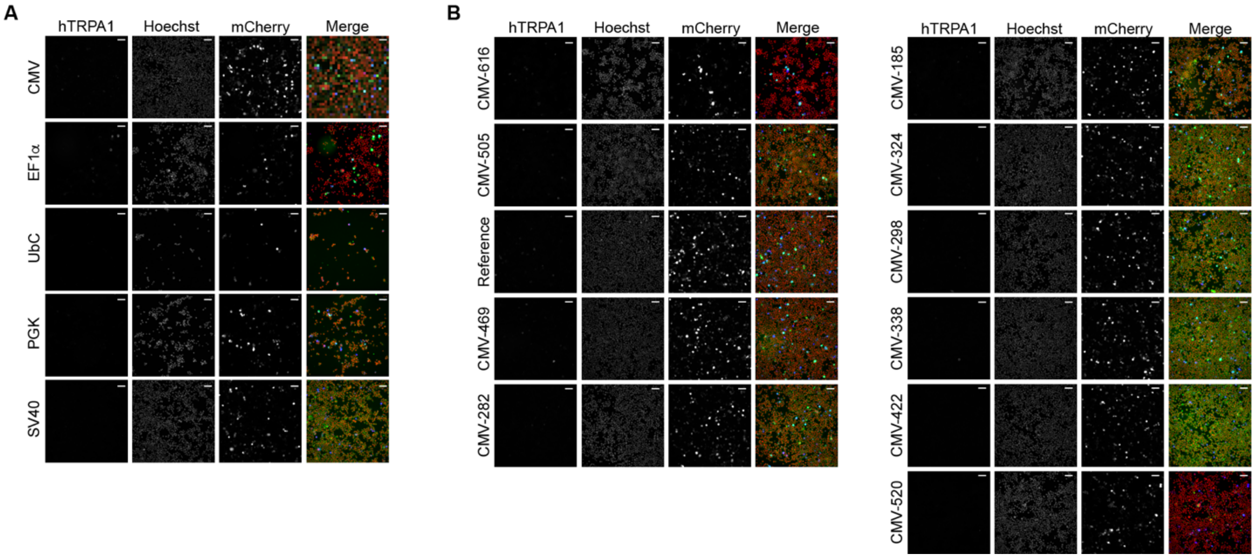


**Figure S1. Representative epifluorescence images of hTRA1 expression in Neuro2A cells controlled by eukaryotic promoters and 5'UTRs.** (**A**) Representative epifluorescence images of Neuro2A cells co-expressing 3xFLAG-hTRPA1 controlled by the indicated eukaryotic promoters and mCherry. Cells were stained with Hoechst (blue) and anti-TRPA1 (green) antibody. Scale bar indicates 64 μm. (**B**) Representative epifluorescence images of Neuro2A cells co-expressing 3xFLAG-hTRPA1 controlled by a CMV promoter without (reference) or with the indicated 5'UTRs and mCherry. Cells were stained with Hoechst (blue) and anti-TRPA1 (green) antibody. Scale bar indicates 64 μm.

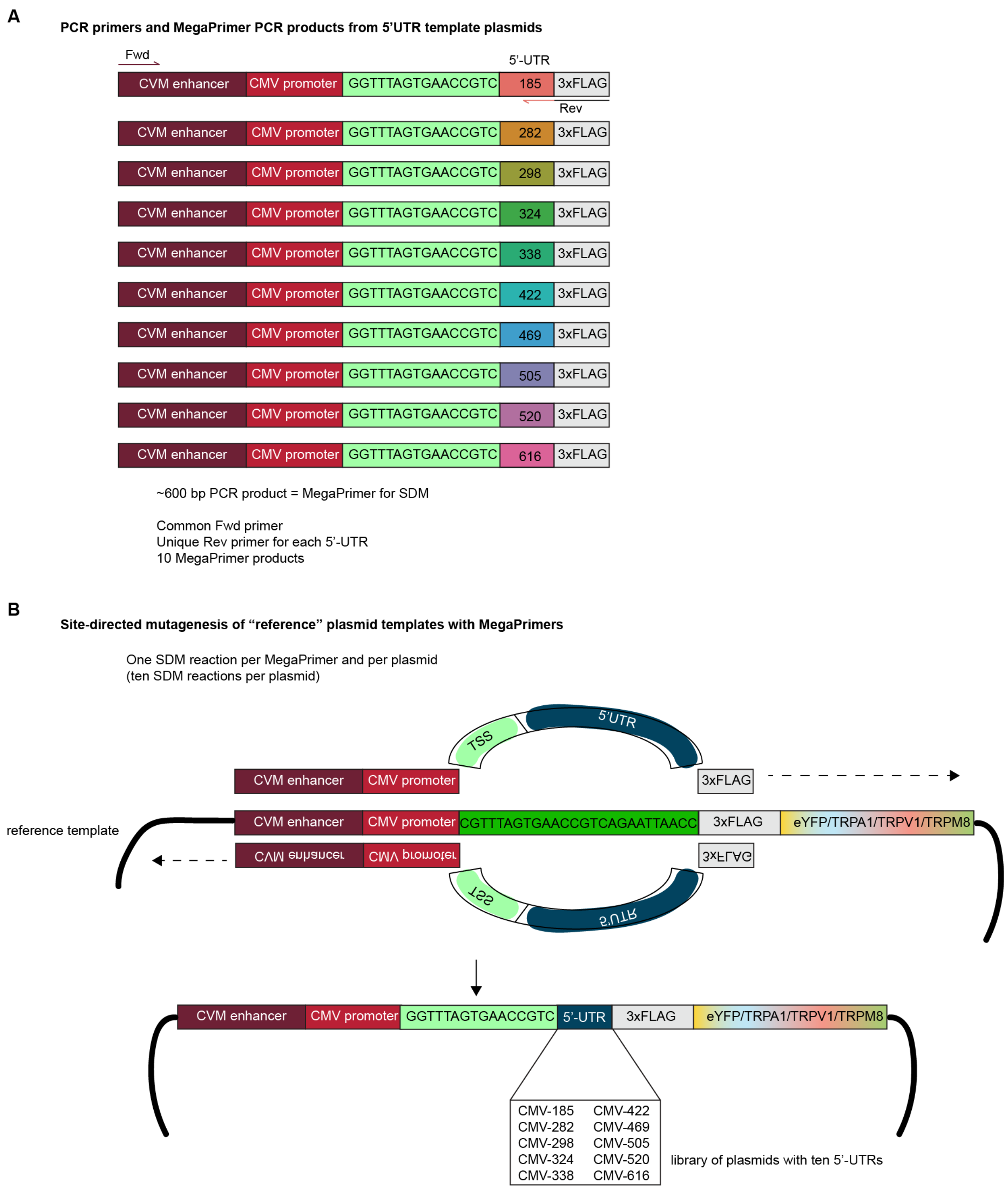


**Figure S2. MegaPrimer-based cloning strategy to easily build library of plasmids with 5'UTRs.** (A) MegaPrimers containing the CMV enhancer (maroon), CMV promoter (red), transcription start site (TSS, light green), 5'UTRs (rainbow colors), and the beginning of the 3xFLAG tag (grey) were generated by PCR amplification from a library of 5'UTR plasmids using a common forward primer (maroon) and unique reverse primers that annealed to the specific

5’UTR and added the beginning of the 3xFLAG tag to allow annealing to target “reference” plasmids. (**B**) These MegaPrimers were then used in a set of site-directed mutagenesis (SDM) reactions to introduce the 5’UTRs into vectors encoding 3xFLAG-tagged proteins. This modular cloning strategy allows the use of one set of MegaPrimers to introduce the 5’UTRs into the p3xFLAG vector independent of the downstream open reading frame (eYFP/TRPA1/TRPV1/TRPM8). The 5’UTR sequences are available in the accompanying Source Data file.

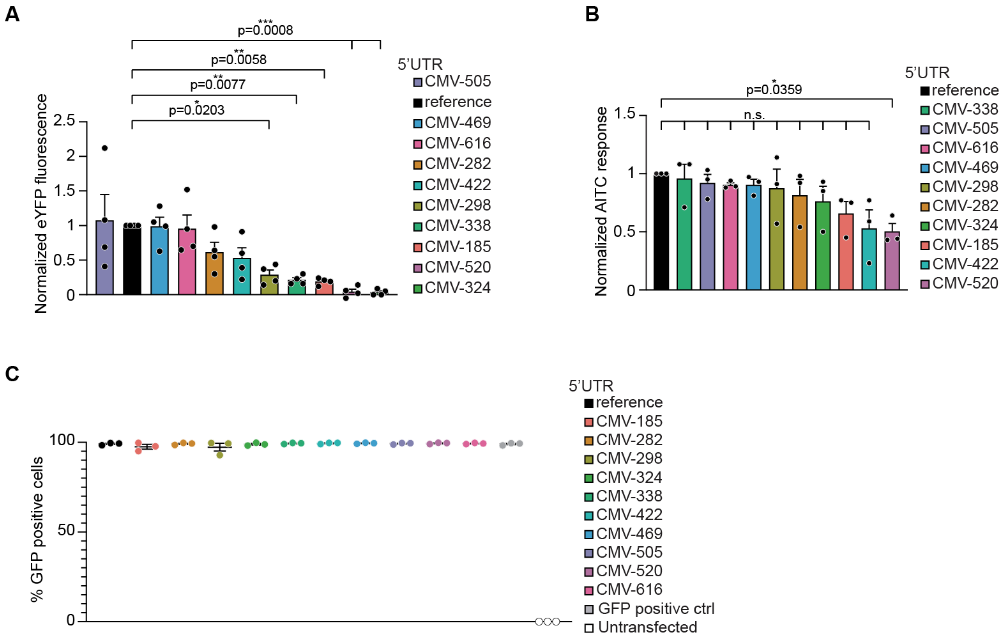


**Figure S3. 5'UTRs provide graded control of eYFP and hTRPA1 expression across cell types.** (**A**) Reference plasmid (CMV promoter, no 5'UTR)-normalized eYFP fluorescence quantification of 3xFLAG-eYFP controlled by the indicated 5'UTRs. Data represent mean ± SEM. $n = 4$ independent experiments, P-values listed in figure panel, one-way ANOVA with Bonferroni's *post hoc* analysis. (**B**) Quantification of reference plasmid-normalized AITC-evoked changes in Fura-2 ratio of 3xFLAG-hTRPA1 controlled by a CMV promoter alone (reference) or with the indicated 5'UTRs from data as in Figure 4A. Data represent mean ± SEM. $n = 3$ independent experiments, with an average of $n \geq 2{,}324$ cells per transfection condition per biological replicate. P-values listed in figure panel, one-way ANOVA with Bonferroni's *post hoc* analysis. (**C**) Quantification of percent GFP positive cells alone (GFP positive control) or that were co-transfected with 3xFLAG-hTRPA1 controlled by a CMV promoter alone (reference) or with the indicated 5'UTR. Untransfected cells were the negative control. Data represent mean ± SEM. $n = 3$ independent experiments, $n \geq 10{,}000$ cells per transfection condition per experiment.

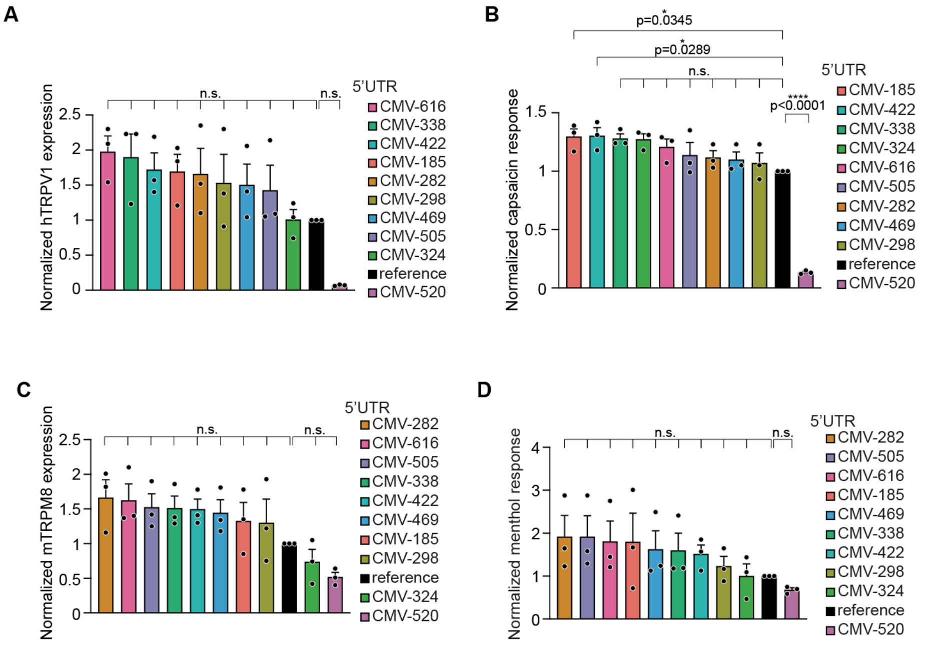


**Figure S4. 5'UTRs predominantly drive high hTRPV1 and mTRPM8 expression.** (**A**) Quantification of the reference plasmid-normalized expression of 3xFLAG-hTRPV1 controlled by a CMV promoter alone (reference) or the indicated 5'UTRs from data as in Figure 6B. Data represent mean ± SEM. $n = 3$ independent experiments, $n \geq 2{,}324$ cells per transfection condition per experiment. P-values listed in figure panel, one-way ANOVA with Bonferroni's *post hoc* analysis. (**B**) Quantification of reference plasmid-normalized capsaicin-evoked change in Fura-2 ratio for 3xFLAG-hTRPV1 controlled by CMV alone (reference) or with the indicated 5'UTRs from data as in Figure 6D. Data represent mean ± SEM. $n = 3$ independent experiments, with an average of $n \geq 2{,}792$ per transfection condition per biological replicate. P-values listed in figure panel, one-way ANOVA with Bonferroni's *post hoc* analysis. (**C**) Quantification of the reference plasmid-normalized expression of 3xFLAG-mTRPM8 controlled by a CMV promoter alone (reference) or with the indicated 5'UTRs from data as in Figure 7B. $n = 3$, one-way ANOVA with Bonferroni's *post hoc* analysis. (**D**) Quantification of reference plasmid-normalized menthol-evoked change in Fura-2 ratio for 3xFLAG-mTRPM8 controlled by CMV alone (reference) or with the indicated 5'UTRs from data as in Figure 7D. Data represent mean ± SEM. $n = 3$ independent experiments, with an average of $n \geq 2{,}724$ per transfection condition per biological replicate. One-way ANOVA with Bonferroni's *post hoc* analysis.

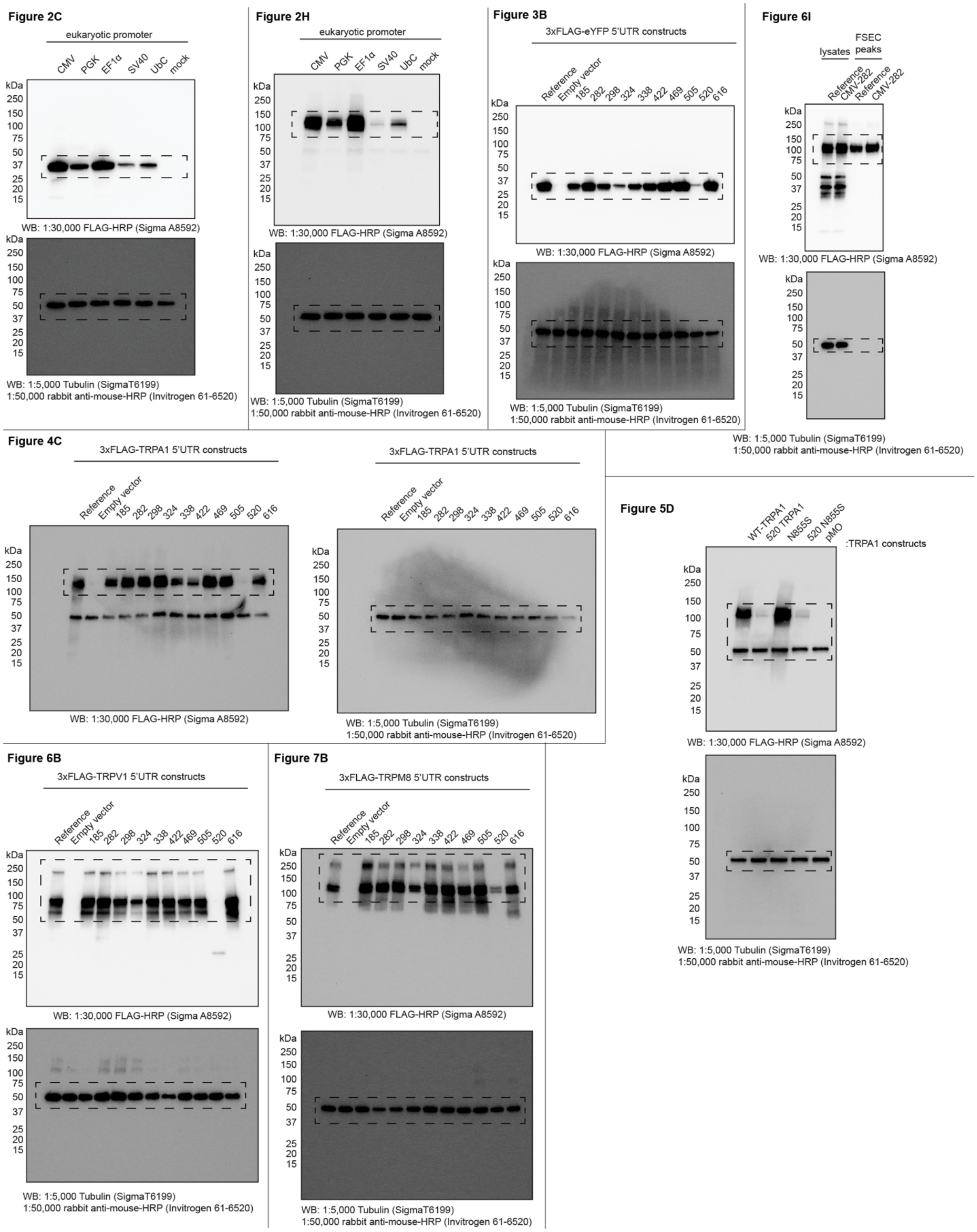


**Figure S5.** Uncut full Western blots shown in Figures 2C, 2H, 3B, 4C, 5D, 6B, 6I, and 7B. The regions surrounded by dashed lines represent the panels in the respective figures.

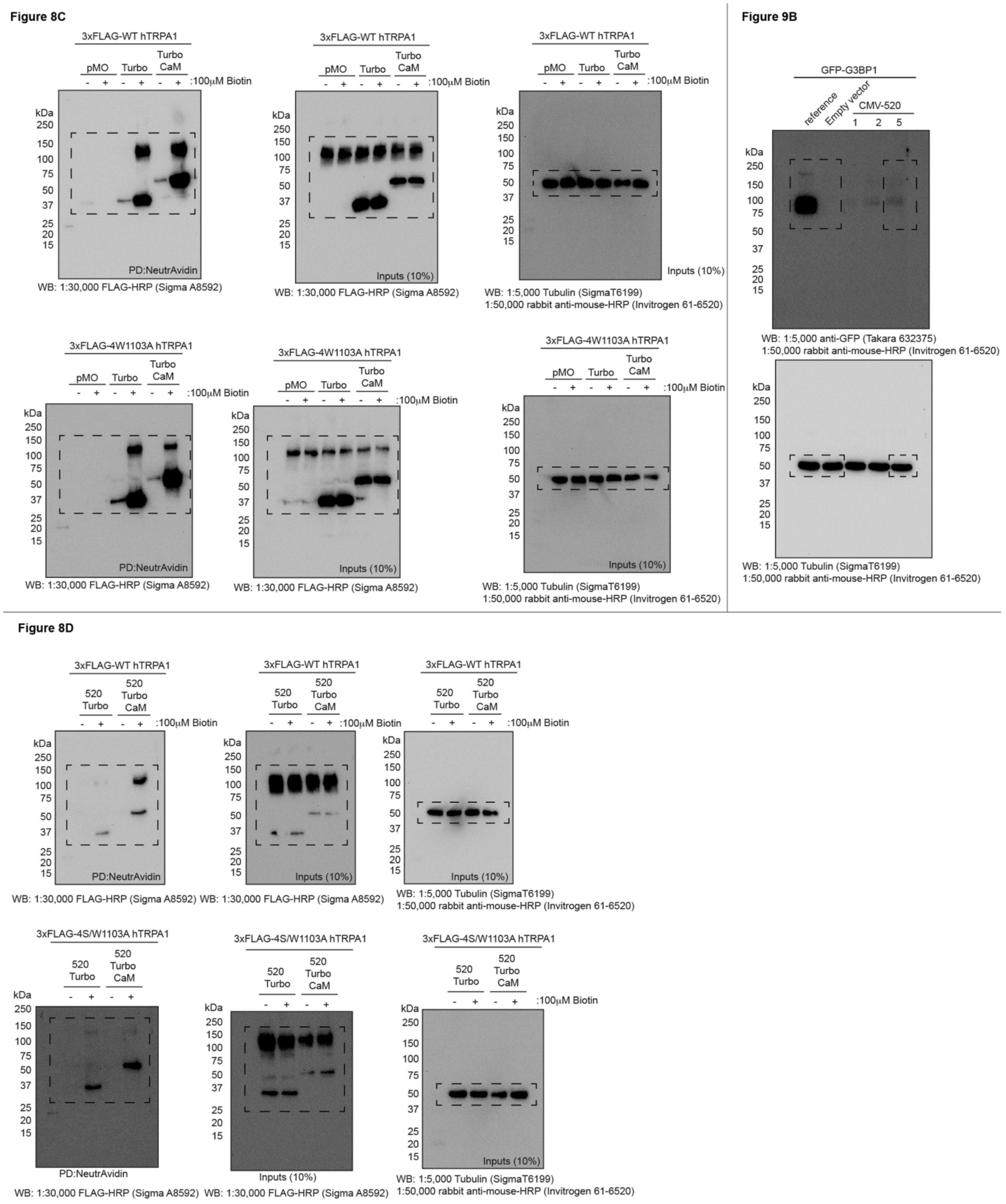


**Figure S6.** Uncut full Western blots shown in Figures 8C, 8D, and 9B. The regions surrounded by dashed lines represent the panels in the respective figures.
